# Unequal mitochondrial segregation promotes asymmetric fates during neurogenesis

**DOI:** 10.1101/2024.11.08.622699

**Authors:** Benjamin Bunel, Rosette Goiame, Xavier Morin, Evelyne Fischer

## Abstract

Asymmetric divisions of vertebrate neural progenitors are critical for generating neurons while preserving the stem cell pool, ensuring proper central nervous system development. It was previously shown that the post-mitotic remodeling of mitochondrial activity in daughter cells influences neural fates. In this study, we demonstrate that unequal distribution of mitochondria during asymmetric mitosis plays a decisive role in triggering this differentiation process. Using live imaging to monitor mitochondrial segregation in individual progenitors and track their progeny’s fate within embryonic neural tissue, we show that daughter cells inheriting fewer mitochondria consistently differentiate into neurons, whereas their sibling receiving more mitochondria retains the progenitor status. Furthermore, experimental displacement of mitochondria during mitosis to force their unequal inheritance was sufficient to drive premature neuronal differentiation. Our findings establish a direct causal relationship between unequal mitochondrial inheritance and the asymmetric fate of sister cells *in vivo*, uncovering a key mechanism in neural development.

## Main text

Asymmetric divisions are a fundamental characteristic of stem cells, relying on the unequal inheritance of fate determinants to drive differentiation of one progeny while simultaneously maintaining stemness in the other daughter. Mitochondria are complex organelles that integrate numerous cellular functions, and their unequal distribution during mitosis would provide an opportunity for the rapid and coordinated modulation of multiple parameters in daughter cells. During vertebrate central nervous system development, the process of neuronal differentiation involves remodeling of mitochondrial functions and metabolism that control downstream gene regulatory pathways (*1*, *2*). Notably, fate decisions in the progeny of cortical progenitors depend on modifications in the balance between mitochondrial fusion and fission taking place within a brief time window following cell division (*2*). This raises the question of what triggers these mitochondrial changes: are they instructed after mitosis downstream of non-mitochondrial fate determinants (*3–5*) or are they independently encoded in an asymmetric mitochondrial behavior within the dividing mother cell?

## Uneven mitochondrial segregation during neurogenesis

We studied mitochondrial segregation during neural progenitor divisions using live imaging in the embryonic chick neural tube. Constructs expressing fluorescent proteins fused to the mitochondrial targeting sequence of Cox8 were electroporated *in ovo* (Figure 1A) and mitochondrial inheritance was monitored *ex ovo* in “en-face” live imaging of the spinal neuroepithelium. This provides optimal access to its apical surface (Figure 1B), where divisions occur. Using 4D confocal microscopy, we monitored mitochondria and cell contours through the whole mitotic sequence at 3min time intervals (Figure 1B, fig.S1A). The mitochondrial volume inherited by each of the two sister cells immediately after cytokinesis was reconstructed and measured. We used a metric termed R_mito_, which represents the ratio of the smaller to the larger of these two measurements, in order to compare mitochondrial inheritance between sister cells (fig.S1B). As an example, a R_mito_ value of 1 indicates that each daughter cell inherits a similar mitochondrial volume (50% of the mother’s mitochondrial pool), whereas a R_mito_ value of 0.5 indicates that one cell inherits twice as many mitochondria as its sister (Figure 1C, fig.S1B). We first monitored mitochondrial inheritance at embryonic day 3 (E3, HH17-18), and found that R_mito_ distributed across a broad range of values between 1 and 0.5 (Figure 1D). At that stage, three modes of progenitor divisions coexist in the spinal cord: symmetric proliferative divisions (producing two progenitors: PP), asymmetric neurogenic divisions (producing a progenitor and a neuron: PN), and a small proportion of symmetric terminal neurogenic divisions (producing two neurons: NN) (*6*, *7*).

**Figure 1:**
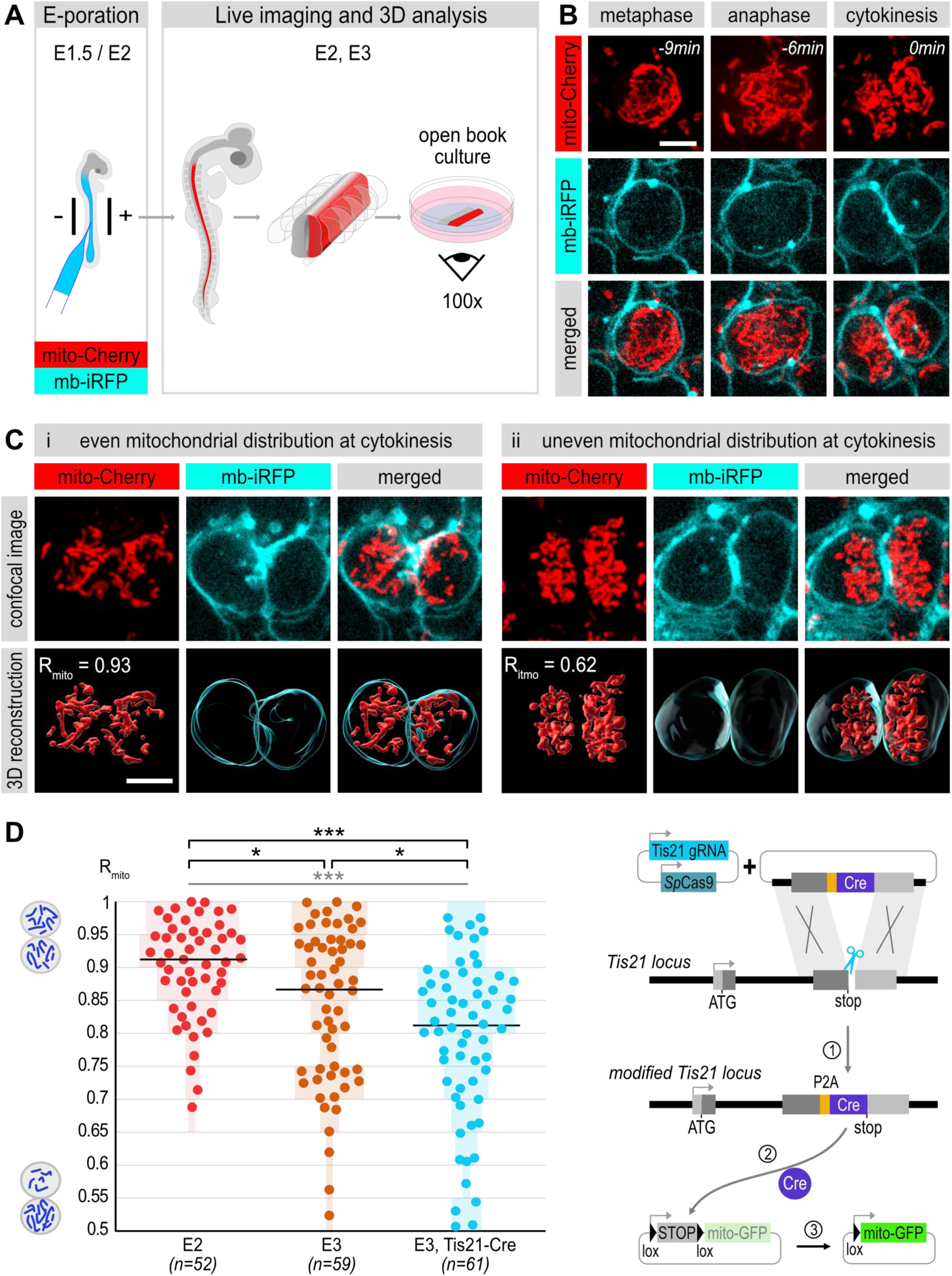
Increased frequency of uneven mitochondrial inheritance at the onset of asymmetric modes of divisions during neurogenesis A. Scheme of *in ovo* electroporation of DNA constructs in the chick embryonic neural tube and experimental strategy for open-book live imaging in the electroporated neural tube. B. En-face time lapse monitoring of mitochondria (red) and cell contours (cyan) in a dividing neural progenitor. Scale bars 5µm. C. En-face confocal view and 3D reconstructions of mitochondrial volume in sister cells immediately after cytokinesis. 3D reconstructed images from the entire z-stacks are used to calculate the ratio R_mito_ of mitochondrial inheritance between sister cells. Typical examples of even (i) and uneven (ii) inheritance are depicted. Scale bars 5µm. D. Left: scatter plots of R_mito_ distribution at E2, E3 and in Tis21 positive progenitors (E3, Tis21- Cre). Light color bars represent the percentage of cell pairs in each 0.05 interval. Statistical analysis: Kruskal-Wallis ***: p=0.0003 (distributions between E2, E3 and E3-Tis21-Cre), Mann-Whitney. ***: p=0.001. *: p=0.026 (E2 vs E3) and *: p=0.015 (E3 vs E3-Tis21-Cre). Right: schematic representation of the CRISPR/Cas9 strategy of Cre recombinase knock-in used to activate a loxP-dependent mitochondrial reporter specifically in Tis21 neurogenic progenitors.

We wondered whether the broad distribution observed at E3 might be linked to the coexistence of these three modes of division. To test this hypothesis, we monitored mitochondrial inheritance at E2, when the symmetric PP mode of division strongly predominates. We found that mitochondria were mostly distributed evenly between sister cells, contrasting with the high frequency of uneven inheritance observed at E3 (Figure 1D). Conversely, we monitored mitochondrial inheritance in a population enriched for progenitors undergoing asymmetric PN divisions at E3. These were identified thanks to a reporter system relying on the targeted insertion of a Cre recombinase at the Tis21 locus, a gene that is turned on at the transition towards neurogenic divisions (*7–10*) (Figure 1D and Materials and Methods). In this subpopulation, the distribution of R_mito_ values shifted significantly towards more unbalanced values compared to the whole progenitor population at E3.

Could the observed differences in mitochondrial volume inheritance be a simple consequence of an overall difference in cell volume between the sisters? To address this question, we monitored both mitochondrial and cellular volumes upon cytokinesis in the daughter cells of Tis21-Cre progenitors. While an important frequency of uneven mitochondrial inheritance was observed in this dataset, the cellular volume was nearly identical between sisters, indicating that these two parameters are uncoupled (fig.S2). These findings confirm that uneven mitochondrial segregation constitutes an independent factor.

## Unequal mitochondrial segregation is a hallmark of asymmetric divisions

The increase in uneven mitochondrial inheritance between pairs of sister cells mirrors the expected rise in the proportion of asymmetric modes of division among the E2, E3 and E3- Tis21 progenitor populations (Figure 1D) (*6–8*). We therefore wondered whether experimental modifying the balance between the different modes of progenitor division would impact the mitotic distribution of mitochondria. We targeted the expression of cell cycle regulators in progenitors, as gain and loss of function for several of these genes have been shown to artificially accelerate or delay the onset of neurogenic modes of divisions (*11–14*). We first experimentally forced premature neurogenic divisions at early stages by overexpressing the CDC25B phosphatase, a G2/M regulator (*11*). This led to a significant shift towards uneven mitochondrial inheritance: the distribution of R_mito_ in CDC25B overexpressing cells at E2.25 was significantly different from the control (Figure 2A), and closely resembled what is normally observed at E3 (see Figure 1D). This higher proportion of uneven mitochondrial inheritance was accompanied by an increased production of neurons 12 hours later (fig.S3), consistent with an increased prevalence of asymmetric modes of division at that stage. Conversely, we knocked-down the G1 regulator CDKN1C/p57^kip2^, as we recently showed that this delays neurogenesis by favoring the symmetric proliferative mode of division (*13*). Remarkably, this significantly reduced the proportion of uneven mitochondrial inheritance at E3 (Figure 2B), and led to a distribution of R_mito_ that closely matched what is normally observed at E2.25 (Figure 2A).

**Figure 2:**
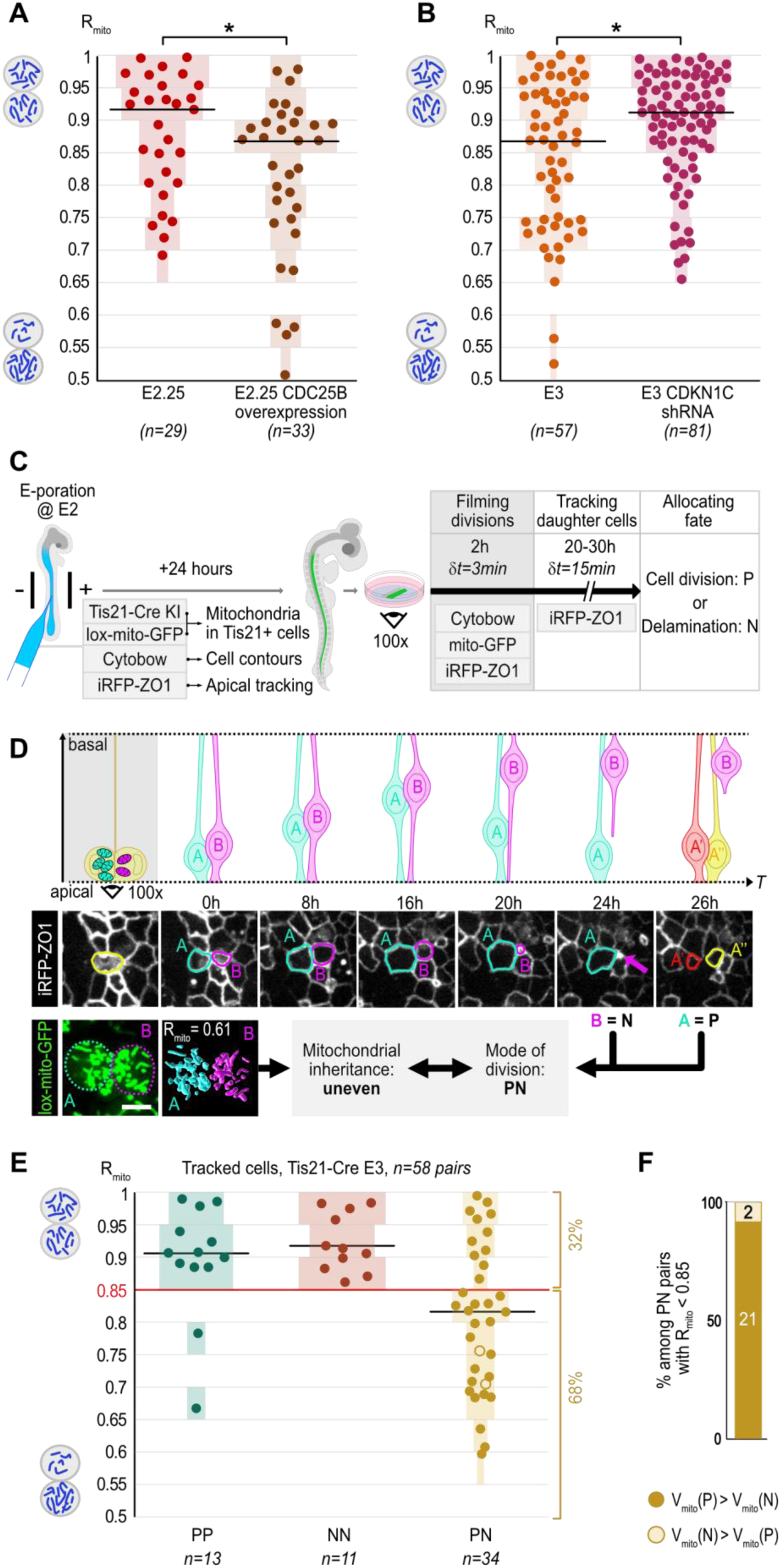
Unequal mitochondrial inheritance by sister cells is a hallmark of asymmetric division A. Scatter plots of R_mito_ in sister pairs of control cells and upon overexpression of CDC25B at E2.25. Mann-Whitney. *: p=0.0199. B. Scatter plots of R_mito_ in control cells and upon downregulation of CDKN1C at E3. Control cell values at E3 are the same as in Figure 1D. Mann-Whitney. *: p=0.019. Light color bars in A and B represent the percentage of pairs in each 0.05 interval. C. Experimental strategy combining live monitoring of mitochondrial inheritance and daughters fate tracking. D. Time-lapse series linking mitochondrial distribution and mode of division in Tis21 positive progenitors. Scheme (top) and en-face view (middle) of the time course from progenitor division to daughters’ fate determination (division: progenitor, delamination: neuron). Bottom: unbalanced mitochondrial inheritance (R_mito_=0.61) matches with an asymmetric PN fate in sister cells. Scale bar: 5µm. E. Scatter plots summarizing R_mito_ and daughter fate (P=progenitor, N=neuron) from Tis21- positive progenitors. Red line: (R_mito_ = 0.85) proposed threshold separating equal from unequal inheritance. F. Inheritance of the smallest mitochondrial pool by the future neuron (full dots in E) or progenitor (open circles in E) in 23 PN pairs with unequal mitochondrial segregation.

To investigate whether uneven partitioning of mitochondria is directly associated with an asymmetric cell fate, we next monitored mitochondrial inheritance in individual progenitors and tracked fate acquisition in their daughter cells through additional imaging of their apical surface for 20 to 30 hours. In the developing spinal cord, which relies on direct neurogenesis, the progressive reduction of the apical surface followed by delamination characterizes a differentiating neuron (N), whereas a new division event identifies a daughter cell as a progenitor (P) (*5*, *15*), and see Materials and Methods; Figure 2C-D and fig.S4).

Our results from 58 pairs of sister cells show that the distribution of R_mito_ differs between symmetrically and asymmetrically dividing progenitors (Figure 2E). In the symmetrically dividing population (PP and NN divisions, n=24 pairs), the values of R_mito_ were homogenously distributed in the 0.85 to 1 range, with the exception of two outliers. We therefore propose a threshold at 0.85 that will be used henceforth to define “equal” versus “unequal” mitochondrial inheritance. In stark contrast to symmetric divisions, the values of R_mito_ in asymmetric divisions were distributed between 0.55 and 1, with the majority of PN daughters (n=23/34) receiving unequal mitochondrial pools (R_mito_ < 0.85) (Figure 2E). This live analysis, which directly links mitotic events in the progenitor to cellular identity post mitosis in its progeny, shows that an unequal mitochondrial inheritance is a strong - albeit not absolute - predictor of an asymmetric fate decision. One additional feature of the individual tracking of pairs of cells is that it enables to discern which daughter cell, of the neuron or the progenitor, receives the largest or smallest mitochondrial pool. We observed a compelling near-exclusive relationship between mitochondrial pool size and fate: in 21 out of the 23 PN divisions showing unequal mitochondrial inheritance, the sister cell receiving fewer mitochondria differentiated into a neuron (Figure 2E and 2F).

Our analysis of pairs of sister cells, and in particular the striking ‘directionality’ of unequal mitochondrial distribution that we observed during asymmetric neurogenic division, further supports the idea that the unequal partitioning of mitochondria during mitosis initiates different mitochondria-regulated programs between progenitors and neurons during embryonic neurogenesis.

## Unequal mitochondrial segregation drives differential fate choices

We then investigated whether the relationship between unequal mitochondrial inheritance and differential fate acquisition is causal. An effective approach involves directly manipulating mitochondrial distribution to induce unequal inheritance in PP progenitors that normally distribute them equally, and subsequently assessing the impact on the identity of their progeny.

### Forcing unequal mitochondrial segregation via artificial attachment to the microtubular network

Previous studies have shown that mitochondrial partitioning during mitosis is essentially symmetric in cultured cell lines, driven by mechanisms including the actin and microtubule networks and their motor proteins (*16–18*). Strikingly, artificially tethering mitochondria to a kinesin to force their attachment to the microtubule network during mitosis disrupts the control of their symmetric distribution to daughter cells (*16*). We developed a time-controlled adaptation of this strategy to force the attachment of mitochondria to the microtubule network during a restricted time window that narrowly encompasses progenitor mitosis (Figure 3A), in order to avoid long-lasting effects on mitochondrial motility in their progeny. We used the CatchFire (chemically assisted tethering of chimera by fluorogenic-induced recognition) method, to induce the rapid and reversible association of mitochondria with a plus-end directed kinesin motor upon application and removal of a specific ligand (*19*).

**Figure 3:**
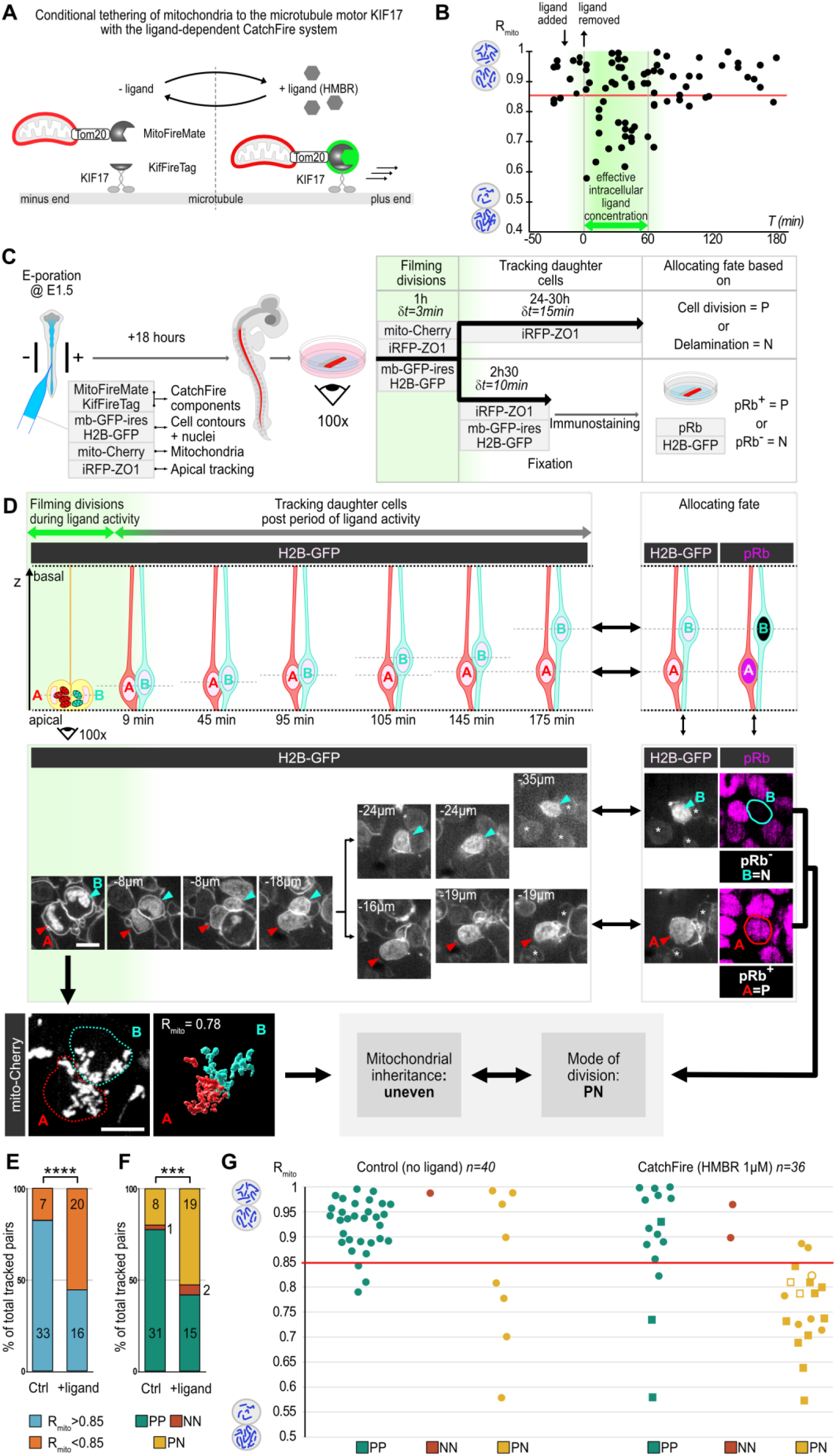
Forced unequal mitochondrial inheritance induces asymmetric fate choices A. Scheme of the CatchFire strategy to force mitochondrial attachment to the microtubular network during mitosis. B. Scatter plots of R_mito_ in CatchFire condition before, with (double arrow) and after washout of ligand. Green box: estimated period of ligand activity in cells. N=80 pairs. C. Combined live monitoring of mitochondrial inheritance and daughter cell fate tracking at E2.5. D. Scheme (top) and en-face views (middle) of daughter cell’s nucleus (H2B-GFP, white) in the depth of the neuroepithelium during the live tracking period (left) and correspondence with pRb immunofluorescence at its end (right, magenta). Bottom: R_mito_=0.78 in the mother cell matches with PN fate: Scale bar: 5µm. Asterisks: neighboring cells used for registration between live and fixed images. **E-G.** Representations of R_mito_ and/or cell fate data from control and CatchFire pairs of cells obtained from the two tracking approaches. E,F: Percentages of R_mito_ < 0.85 vs > 0.85 (E) and fate (F). Numbers in columns represent the number of pairs. G: Scatter plots summarizing R_mito_ and daughter fate data. Squares in the CatchFire condition indicate a mitochondrial compaction index (Ci)< 0.6. Open symbols in PN pairs: smallest pool inherited by progenitor. P=progenitor, N=neuron. Chi2 test: E, p=0.0005 ; F, p=0.006.

We electroporated vectors expressing fusions of the CatchFire dimer moieties, one targeted to the outer mitochondrial surface (MitoFireMate) and the other fused to a kinesin (KifFireTag), (Figure 3A) together with fluorescent mitochondrial and membrane reporters, and assessed mitochondrial behavior following administration of the ligand in en-face live preparations. In mitotic cells, within 15 minutes of ligand exposure, mitochondria located more basally than in controls and appeared more compacted from metaphase until cytokinesis (fig.S5 and fig.S6). Most importantly, this led to unequal mitochondrial segregation, including extreme R_mito_ values lower than 0.5 (fig.S6B), demonstrating that the CatchFire approach reproduces *in vivo* what has been observed upon artificial tethering of mitochondria to a kinesin *in vitro* (*16*). We then calibrated the ligand administration regime to obtain R_mito_ values contained within the physiological range observed in asymmetrically dividing progenitors, and to restrict the effect of the ligand to the approximate duration of mitosis (see Materials and Methods for details). With this optimized protocol, we monitored mitotic events at E2, when a majority of progenitors undergo PP divisions and mitochondrial inheritance is mostly equal (Figure 1D). We observed a significant shift towards unequal inheritance in ligand-treated cells, whereas the great majority of cells dividing before and after ligand exposure displayed the characteristic equal mitochondrial segregation normally observed at that stage (Figure 3B and fig.S6C; see also Figure 1D). Of note, cells that retained a R_mito_ value higher than 0.85 during ligand exposure did not display any mitochondrial compaction (fig.S6D-F), suggesting that these cells did not receive the two components of the CatchFire system, as can happen when multiple vectors are co-electroporated (*20*).

The range of experimentally-induced R_mito_ values obtained at E2 with this molecular toolkit is comparable to what is normally observed at E3 (Figure 1D) and should not compromise daughter cell survival. Hence, the powerful CatchFire approach that we have developed offers a unique opportunity to unravel the functional link between mitochondrial distribution and cell fate decisions.

### Forcing unequal mitochondrial segregation is sufficient to induce asymmetric fate choices

In the next series of experiments, we combined CatchFire manipulation of mitochondrial distribution with daughter cell fate tracking. We first monitored mitochondrial distribution in progenitors undergoing mitosis during the 1 hour period of CatchFire ligand activity. We then used two complementary approaches to establish cell fates (Figure 3C). In one of them, we tracked daughter cells for 24 to 30 hours, allowing identity to be assigned from a new round of mitosis or an apical detachment, as performed previously (Figure 2C-D). In the other set of tracking experiments, we imaged daughter cells for only 2.5 additional hours, then fixed the samples and used molecular characterization to assign an identity. We took advantage of the differential expression of the hyperphosphorylated form of the Retinoblastoma protein (pRb), which is detected in all cycling progenitors (see Material and Methods), whereas future neurons remain negative for this marker (*13*). Daughter cells positive or negative for pRb at the end of the 2.5 hours tracking period could therefore be identified as a cycling progenitor (P) or a neuron (N), respectively (Figure 3D and fig.S7).

In these experiments, the increase in the number of pairs showing unequal mitochondrial segregation in the CatchFire condition (Figure 3E, fig.S8A) matched with a rise in the proportion of asymmetric PN divisions, with more than half of the pairs (53%, n=19/36) showing one sister cell committed to differentiation, compared to only 20% (n=8/40) in the control condition (Figure 3F, fig.S8B). This premature shift toward a neuronal fate in one of the daughter cells was observed specifically in pairs of sister cells displaying unequal mitochondrial inheritance (Figure 3G, fig.S8C).

The shift was even more compelling when focusing on the 15 pairs with CatchFire-induced increased mitochondrial compaction (squares in Figure 3G): 14 of these pairs displayed unequal mitochondrial inheritance, and we observed a different sister fate (PN) in 12 of them (86%, n=12/14; Figure 3G). Additionally, the relationship between mitochondrial volume inheritance and fate determination was conserved: the smaller mitochondrial pool was inherited by the future neuron in 10 of these 12 PN pairs (Figure 3G), consistent with our observations in asymmetrically dividing progenitors at E3 (Figure 2E).

Overall, these results show that an experimentally induced mitochondrial imbalance during division is sufficient to induce a neurogenic fate in the daughter cell that inherits fewer mitochondria.

Our *in vivo* data show that during embryonic neurogenesis, a quantitative imbalance in the distribution of mitochondria during mitosis is sufficient to establish a new identity in the daughter cell receiving the smaller pool. These data expand upon previous *in vitro* studies in non-neural adult stem cells which have shown a correlation between inheritance of qualitatively distinct mitochondrial pools and different daughter cell proliferative capacities, and introduce the notion of a role for mitochondrial volume inheritance in fate decisions (*21–25*). This raises the intriguing possibility that quantitative apportioning is a distinctive feature of the developmental context, characterized by rapid tissue growth over a short timescale and sustained production of organelles. This may be different in the comparatively much slower process of adult homeostasis, which are more prone to the accumulation of damaged proteins and organelles, and in which the qualitative sorting and exclusion of aging or dysfunctional mitochondria could be necessary for the long-term preservation of the stem cell pool.

How could a smaller mitochondrial volume cause a fate switch towards differentiation? One possibility is that inheriting fewer mitochondria causes an energetic or metabolic deficiency after mitosis. This may influence the post-mitotic remodeling of the mitochondrial network, and in turn influence cell fate. Such an hypothesis is consistent with previous data showing that inhibiting fission prevents differentiation (*1*, *2*), whereas fission favors an increased OXPHOS metabolism which promotes neural differentiation (*1*, *2*) and concomitantly represses progenitor pathways (*26*, *27*). In parallel, this initial signal may trigger the progressive increase of mitochondrial mass that is observed later during differentiation (*28*). This mechanism, whereby the differentiation-specific metabolic shift is primed by a reduced mitochondrial volume at mitotic exit, may represent a new pathway linking these organelles to cellular fate decisions in a developmentally regulated manner.

Our live imaging and fate tracking experiments present the first direct *in vivo* evidence that mitochondria act as asymmetric cell fate determinants in neural progenitors. Deciphering whether and how mitochondrial segregation is coordinated with the mitotic distribution of other determinants involved in asymmetric fate decisions in spinal progenitors (*3–5*), and exploring the interplay between mitochondria and their downstream pathways, will represent challenges for future research.

## Supporting information

Bunel et al - Movie S4

Bunel et al - Movie S5

Bunel et al - Movie S6

Bunel et al - Movie S7

Bunel et al - Movie S8

Bunel et al - Movie S1

Bunel et al - Movie S2

Bunel et al - Movie S3

## Acknowledgements

We thank Samuel Tozer for insightful discussions on the project, and our colleagues Jean-François Brunet, Sonia Garel, Alice Meunier, Jonathan Weitzman, Marco Pontoglio and Samuel Tozer for their comments on the manuscript. We thank Fabienne Pituello for the kind gift of CDC25B plasmids, and Octave Joliot, Franck Perez and Arnaud Gautier for generously sharing unpublished components of the CatchFire system. We thank the IBENS imaging platform for access to the IMARIS analysis station. **Funding**: work in the Morin lab was supported by grants from the Agence Nationale de la Recherche (SYMASYM ANR-18-CE16- 0021-01), the Fondation pour la Recherche Médicale (FRM EQU202003010547) and the Labex MEMOLIFE. This work has received support under the program « Investissements d’Avenir » launched by the French Government and implemented by the ANR, with the reference ANR- 10-LABX-54 MEMO LIFE. **Authors contributions:** Conceptualization: E.F.; Formal analysis: B.B., E.F.; Funding acquisition: X.M.; Investigation: B.B., R.G., X.M., E.F.; Methodology: B.B., X.M., E.F.; Project administration: X.M., E.F.; Supervision: E.F.; Validation: E.F.; Visualization: B.B., X.M.; Writing – original draft: E.F.; Writing – review & editing: X.M. and E.F.

## Data and materials availability

All data are available in the manuscript or the supplementary materials. Plasmids used in this study and the full sequences are available upon request to X.M.

## Materials and methods Embryos

JA57 chicken fertilized eggs were provided by EARL Morizeau (8 rue du Moulin, 28190 Dangers, France). They were incubated at 38 °C in a Sanyo MIR-253 incubator for the appropriate time. The sex of the embryos was not determined. Under current European Union regulations, experiments on avian embryos between 2 and 4 days *in ovo* are not subject to restrictions.

### *In ovo* electroporation

*In ovo* electroporations in the chick neural tube were performed at embryonic day 1.5 (E1.5, HH stage 10) or E2 (E2, HH stage 14). The DNA solution, containing the vital dye FastGreen for visualization, was injected directly into the lumen of the neural tube via glass capillaries. Five 50ms pulses of 20V (at E1.5) or 25V (at E2) separated by 100 ms intervals were applied using a square-wave electroporator (Nepa Gene, CUY21SC) and a pair of 5 mm gold-plated electrodes (BTW Genetrode model 512) separated by a 4 mm interval. The egg was then sealed with parafilm and returned to the incubator.

### Plasmids

A list of plasmids used in this study is presented in Table S1

#### Mitochondrial reporters

The Mitobow construct (X503) has been described in (*29*). pCX-lox-H2B-EBFP2-lox-mitoGFP (X1017, lox-mito-GFP in Figures) was created as follows: a pCX-lox-H2B-EBFP2-lox-mitoCerulean vector (X992) was first generated by deletion of a SacI-PmlI fragment from Mitobow construct. A lox-H2B-EBFP2-polyA-lox-mito PCR fragment was then amplified from X992 with the following oligonucleotides (CX-fw 5’- GTCTCATCATTTTGGCAAAGAATTCTAGCT-3’ and GFP-Eco-mito-rev 5’- CCTTGCTCACCATGGTGGCGAATTCGGAGGTGGCGACGG-3’) and Gibson cloned into NheI/AgeI digested X-013 (pCX-GFP, (*30*)).

pCX-mito-Cherry (X1202, mito-Cherry in Figures) was created as follows: a pCX-lox-mito-Cherry-lox-mito-Cerulean vector (X993) was first generated by deletion of a SmaI-PacI fragment from Mitobow construct. A lox-mito-Cherry-polyA-lox-mito PCR fragment was then amplified from X993 with CX-fw and GFP-Eco-mito-rev oligonucleotides and Gibson cloned into NheI/AgeI digested X-013 (pCX-GFP, (*30*)) to construct X-1118. pCX-mito-Cherry (X1202) was obtained by removing the BsrGI (stop-lox-mito-GFP) fragment from X-1118.

#### CatchFire plasmids

pK-Tom20-FireMate and pK-Kif17a-FireTag (unpublished vectors related to (*19*), generously provided by Arnaud Gautier, Frank Perez and Octave Joliot) were modified for optimized expression in the chick neural tube by replacing the CMV promoter with a CAGGS (CX) promoter. To generate pCX-Tom20-FireMate (MitoFireMate in Figures), the CMV promoter in pK-Tom20-FireMate was replaced by the Nde1/Nhe1 CAGGS promoter fragment from pCX-MCS2 (*30*). To generate pCX-Kif17a-FireTag (KifFireTag in Figures), the CMV promoter in pK-Kif17a-FireTag was replaced by a Nde1/Mfe1 fragment from pCX-H2B- miRFP670.

#### Membrane contours

pCX-mb-GFP-ires-H2B-GFP, pCX-mb-iRFP670 (cell membranes) or Cytobow (cytoplasm).

#### Apical staining

pCX-ZO1-miRFP670.

#### Challenge of mode of divisions

PMP-CDC25B, PMP-lacZ were kind gifts from Fabienne Pituello’s Lab. The shCDKN1C vector (X1101, (*13*) contains a chick CDKN1C miRNA under the control of chick U6 promoter, H2BGFP under control of CAGGS promoter.

#### Somatic knock-in of the Cre recombinase at the Tis21 locus

Somatic knock-in of the Cre recombinase coding sequence at the C-terminus of the Tis21 locus was achieved via CRISPR-Cas9-based Homology-Directed Recombination (HDR), through co-electroporation of a donor vector (Tis21-P2A-Cre, X907) and a CRISPR/Cas9 vector expressing the Cas9 recombinase and a gRNA targeting the C-terminus of the Tis21 locus (pCX-SpCas9-cTis21-gRNA2, X854). The Tis21 gRNA targets the cttggcagtgtaaggacaag sequence on the antisense strand and cuts 11 bases downstream of the Tis21 stop codon. It was cloned in the BpiI-linearized X330 vector as described (*31*) using a duplex of oligonucleotides Tis21-gRNA2-fw (5’-CACCGCTTGGCAGTGTAAGGACAAG-3’) and Tis21-gRNA2-rev (5’-AAACCTTGTCCTTACACTGCCAAGC-3’). The Tis21-P2A-Cre donor vector consists of a P2A-Cre cassette flanked with long left (1010bp) and right (971bp) arms of homology to the C-terminal region of chick Tis21 (bGalGal1.mat.broiler.GRCg7b genome assembly). The P2A pseudo-cleavage sequence ensures that the Tis21 protein and Cre recombinase are produced as two independent proteins. In order to prevent targeting of the knock-in plasmid and re-targeting of the locus after insertion of the knock-in cassette by the gRNA, the gRNA target sequence was destroyed in the right arm of homology via the introduction of 7 bases changes. Full detail of the construction of this vector and of its validation as a bona fide Tis21 targeting vector are described in (*8*).

### Plasmid concentrations for monitoring the mitotic mitochondrial distribution at different stages of neurogenesis and functional challenges

For E2 experiments, pCX-mb-GFP-ires-H2B-GFP (0.5µg/µl) and pCX-mito-Cherry (0.3µg/µl) were electroporated at E1.5.

For E3 experiments, either pCX-mito-CFP (0.5µg/µl) or a mix of Mitobow (0.5µg/µl) and pCX- Cre (0.1µg/µl) were electroporated with pCX-mb-iRFP670 (0.3µg/µl) at E2.

For measurements in Tis21 positive progenitors at E3, Tis21-P2A-Cre donor and pCX-SpCas9- cTis21-gRNA2 vectors were each used at 0.8 µg/µl in the electroporation mix. The pCX-lox- H2B-EBFP2-lox-mito-GFP reporter plasmid (0.3µg/µl) and pCX-mb-iRFP670 (0.5µg/µl) were added to the electroporation mix.

For experiments forcing premature neurogenic divisions at E2.25, pCX-mb-GFP-ires-H2B- GFP (0.5µg/µl), pCX-mito-Cherry (0.3µg/µl) and either PMP-CDC25B (1µg/µl) or the control vector PMP-lacZ (1µg/µl), were electroporated at 1.5.

For experiments forcing symmetric proliferative divisions at E3, pCX-mb-iRFP670 (0.5µg/µl), pCX-mito-Cherry (0.3µg/µl) and shCDKN1C (1µg/µl) were electroporated at E2.

### Plasmid concentrations for neurogenesis rate measurements

E1.5 embryos were electroporated with pCX-H2B-GFP (0.3µg/µl) and either PMP-CDC25B (1µg/µl) or PMP-lacZ (1µg/µl) for controls and they were harvested 30 hours after electroporation.

### Plasmid concentration for fate tracking in Tis21+ progenitors

E2 embryos were electroporated with Cytobow (0.3µg/µl), pCX-lox-H2B-EBFP2-lox-mito-GFP (0.3µg/µl), pCX-ZO1-miRFP670 (0.3_µg/µl), Tis21-P2A-Cre (0.8µg/µl) and pCX- SpCas9-cTis21-gRNA2 (0.8µg/µl).

### Plasmid concentration for CatchFire experiments

E1.5 embryos were electroporated with pCX-Tom20-FireMate (0.3µg/µl), pCX-Kif17-FireTag (0.3µg/µl), pCX-mb-GFP-ires-H2B-GFP (0.4µg/µl), pCX-mito-Cherry (0.4µg/µl) and pCX- ZO1-miRFP670 (0.4µg/µl).

### Imaging

All the images in this study were obtained on an inverted microscope (Nikon Ti Eclipse) equipped with a spinning disk confocal head (Yokogawa CSUW1) with Borealis system (Andor) and a sCMOS Camera (Orca Flash4LT, Hamamatsu) and the following Nikon objectives: 10x (CFI Plan APO λ, 0.45), 40x (CFI Plan APO λ, 0.95) or 100x (CFI APO VC, NA 1.4, oil immersion). The setup is driven by MicroManager software (*32*) and equipped with a heating enclosure (DigitalPixel, UK) set to 38°C for live imaging experiments.

### En-face culture

En-face culture of the embryonic neuroepithelium was performed at E2.25 or E3. After extraction from the egg and removal of extraembryonic membranes in 1xPBS (thereafter PBS), embryos were transferred to 38 °C F12 medium and pinned down with dissection needles at the level of the hindbrain in a 35 mm Sylgard dissection dish. A dissection needle was used to slit the roof plate and separate the neural tube from the somites from hindbrain to caudal end on both sides of the embryo. The neural tube and notochord were then transferred in a drop of F12 medium to a glass-bottom culture dish (MatTek, P35G-0-14-C) and medium was replaced with 1ml of 1% low melting point (LMP) agarose/F12 medium (maintained at 38°C). Excess medium was removed so that the neural tube would flatten with its apical surface facing the bottom of the dish, in an inverted open book conformation. After 30 seconds of polymerization on ice, an extra layer of agarose medium (200 µl) was added to cover the whole tissue and left to harden. 2ml of 38 °C culture medium was added (F12/1mM Sodium pyruvate) and the culture dish was transferred to the 38 °C chamber of a spinning disk confocal microscope.

### Live monitoring of mitochondria during cell divisions

Several full-frame fields of view (136*136µm) were selected on the flat mounted neural tube, and ∼25µm deep z-stacks were acquired with a 0.3µm z-step size at 3 minutes intervals during 1 to 2 hours. Images were saved as separated images and imported in Fiji. Pairs of daughter cells at the first timepoint after division presenting both membrane or cytoplasmic reporter signal and mitochondrial signal were identified and selected to be cropped on their entire height. Selected crops were saved as image stack files for 3D reconstruction (see below).

### Daughter fate tracking experiments

For fate tracking experiments, an initial acquisition of images was performed using the parameters described above for mitochondrial monitoring for a duration of 1 to 2 hours, before switching to new acquisition parameters as detailed below.

Of note, mitochondrial ratio and daughter fates were determined independently in a blind manner within the whole set of pairs. Mode of division and mitochondrial inheritance were matched retrospectively.

For long term live tracking, the acquisition parameters were modified as follows: ∼25µm deep z-stacks centered around the apical surface were acquired with a 1µm z-step at 15 minutes intervals for 24 to 30 hours. To reduce potential phototoxicity, the mitochondrial channel was omitted during this sequence as it is not required for tracking. Using the apical surface marker (ZO1-miRFP670), we were then able to track daughter cells and assign their identity from their behavior: progressive apical constriction (over at least three consecutive time points) followed by apical detachment for a future neuron, versus apical surface division for a progenitor. The purpose of keeping a large z-depth centered on the apical surface was to accommodate for potential z-drifting of the embryo relative to the coverslip during the long acquisition.

For tracking experiments combining short-term live tracking and immunostaining, we used the following protocol: we increased the depth of imaging to 50µm, and acquired 1µm-spaced z-stacks every 10 or 15 minutes for 2h to 2h30. We then switched to the 10x objective and acquired an extended field of view to acquire spatial landmarks in order to facilitate the repositioning of the culture dish with the same orientation after immunostaining. The culture dish was then removed from the microscope stage and immediately processed for fixation and immunostaining. The whole staining procedure was performed at 4°C, and we only subjected the samples to very gentle agitation to ensure that the neural tubes remained embedded in the 1% LMP agarose layer. The culture medium was removed, and the samples were quickly washed once in PBS before fixation for 90min in ice-cold 4% paraformaldehyde (PFA) in PBS followed by 3 washes of 15 minutes in PBS. The neural tube was incubated for 48h with the anti-pRb and anti-GFP primary antibodies in the blocking solution (PBS/0,3% Triton/10%Fetal Calf Serum (FCS)). The samples were then washed 3 times for 15 minutes in PBS and incubated for 48 hours in the dark with the secondary A488 anti-chicken (1/500) and Cy5 anti-rabbit (1/500) antibodies and DAPI (1/1000) in PBS/0,3% Triton. The samples were then washed 3 times for 15 minutes in PBS. 1 ml of PBS was added to the culture dish, which was subsequently transferred to the microscope. We first used the 10x objective to reposition the dish in the same orientation and recover each original field thanks to the 10x images acquired at the end of the 2h30 movie. We then switched to the 100x objective, and used the last images of the 100x live sequence to precisely recover the corresponding fields and acquire matching 4 channels z-stacks in the immune-stained neural tube (405nm/DAPI; 490nm/mb-GFP and H2B-GFP; 561nm/mito-Cherry; 641nm/Cy5-pRb).

We then used the H2B-GFP signal in the live sequence to track daughter cells as their nucleus starts to move basally after mitosis. Briefly, for each field of view, we imported the stack files of the 1 hour and 2h30 live sequences in Fiji. We identified divisions of electroporated progenitors occurring during the high resolution 1 hour movie and we followed the nucleus of the daughters during the 2h30 tracking movie. For each pair in which we were still able to visualize the nucleus in each sister cell at the last time point, we proceeded to analyze the corresponding 100x z-stack from the fixed sample and recovered these sister cells on the basis of their H2B-GFP signal. We then determined the identity of the two daughter cells on the basis of the pRb immunostaining in the far-red channel (pRb (+) = progenitor; pRb (-) = neuron).

The 2.5-hour time window was established beforehand in E2 embryos as follows: we determined the time point after mitosis at which all progenitors in a synchronized cohort of cells undergoing mitosis have crossed the restriction point/late G1 stage, as determined by pRb immunoreactivity. We used the cell-permeant dye FlashTag (FT) to label a cohort of pairs of sister cells that perform their division synchronously at E2, and counted the proportion of pRb-positive cells in the cohort in transverse sections from embryos harvested at different time points after FT injection. This proportion should reach a plateau when all the progenitors in the cohort have passed the restriction point and have become positive for pRb. At E2, this plateau was reached between 1.5 hours and 2 hours after injection, and the distribution of pRb immunoreactivity in FT-positive cells was stable at later time points (2.5 hours and 3 hours). This indicates that from 2 hours after mitosis injection, pRb-positivity is a reliable marker of progenitor status, and that a pair of cells containing 0, 1, or 2 pRb-positive cells correspond to a NN, PN and PP pair, respectively. For these experiments, a 1mM stock solution of FlashTag was prepared by adding 20μl of DMSO to a CellTrace Far red dye stock vial (Life Technologies, #C34564) as previously described (*33*). A working solution of 100μM was subsequently prepared by diluting 1μl of stock solution in 9μl of 38°C pre-heated PBS, and injected directly into E2 chick neural tubes. The eggs were resealed with parafilm and embryos were incubated at 38 °C for the appropriate time until dissection.

### 3D reconstruction and measurement of cellular and mitochondrial volumes

Images of pairs of daughter cells at the first time point after division were imported as stacks in IMARIS software (Bitplane) to perform 3D reconstructions of cell volume and mitochondrial volume. Each daughter cell was individually segmented and reconstructed. Cells were delineated by drawing their contour on each z plane through their entire height using either the membrane reporter signal or a cytoplasmic reporter signal. With the contouring of the daughter cells, the cellular volume of each daughter cell was reconstructed using the surface creation tool of IMARIS. The mitochondrial signal contained in the cellular volume of each daughter cell was then isolated by masking mitochondrial signal located outside of this volume. Each daughter’s mitochondrial volume was then reconstructed with the surface creation tool, using the same threshold for both cells. Data of both mitochondrial and cellular volumes for each daughter cells were then used to perform ratio measurements to assess the mitochondrial distribution between daughter cells and to compare their cellular volume.

### Forced attachment of mitochondria to kinesin using CatchFire

We developed a time-controlled strategy to reversibly force mitochondrial attachment to the microtubular network (Figure 3A). The rationale was to restrict the perturbation to a time window around progenitor division in order to avoid long lasting effects on mitochondrial motility that may perturb their activity and be deleterious for daughter cell survival. The recently developed CatchFire (chemically assisted tethering of chimera by fluorogenic-induced recognition) method is based on the reversible ligand-dependent dimerization of small genetically encoded protein partners. The smaller moiety of the dimer (FireTag) was fused to the N-terminus of the motor domain of the plus-end kinesin-like KIF17 (*19*) (thereafter called KifFireTag). The larger moiety (FireMate) was anchored to mitochondria via the targeting sequence of the outer mitochondrial membrane protein Tom 20 (MitoFireMate). The two partners can therefore enable the rapid and reversible association of mitochondria with the microtubule motor upon ligand administration and removal, respectively (Figure 3A).

Embryos were electroporated at E1.5 with the appropriate plasmid combination (see above) and processed 18 hours later for en-face live-imaging according to the procedure described above. To induce the pairing of KifFireTag and MitoFireMate, a 20µM solution of ligand HMBR was prepared by diluting the original 20 mM stocks in F12 medium, and an appropriate volume was added to the culture dish to reach the desired final concentration of ligand. We first established a protocol of transient exposure of the sample to ligand-containing medium, that considers the diffusion kinetics of the ligand within the complex neuroepithelial tissue. Based on our previous experience with the HMBR ligand and pFAST, the fluorogenic monomeric parent of the CatchFire system, we expected that the diffusion of the ligand within the tissue would take between 15 and 30 minutes, and that it would display a slow disappearance kinetics after being washed out from the culture dish (*34*). We therefore anticipated that the induction and cessation of pairing of the CatchFire partners would follow a similar kinetics. We found that continuous exposure (1 hour) and/or high ligand concentration (10µM) led to aberrant mitochondrial distribution (fig.S6B) and mitosis failure in several dividing cells. After several tests, we determined an optimal HMBR concentration (1µM) and duration of exposure (15minutes) after which ligand removal from the culture medium initiates a period of progressive depletion of the ligand from neuroepithelial cells. With this protocol, the intracellular ligand concentration appears to remain in the range of efficient CatchFire pairing for approximately 1 hour before the effect disappears (as judged from measurements of mitochondrial inheritance and compaction index - see Figure 3B and fig.S6E). In the final protocol used for fate tracking experiments, we started by selecting several fields of interest for imaging in the absence of the ligand. We then added ligand to the dish to reach a 1µM concentration in the medium. After 15 minutes, the ligand was removed from the culture dish by three gentle washes with 2 mL of HMBR-free F12 medium. We immediately started imaging progenitor divisions in the selected fields for 1 hour using the acquisition parameters described above in the “Live monitoring of mitochondria during cell divisions” section. After the 1-hour movie, we switched to the relevant acquisition parameters describe above in the “Daughter fate tracking experiments” section.

For the live visualization of the microtubule network (fig.S5 and fig.S6A) dissected neural tubes were incubated in a SiR-tubulin solution (SC006: Cytoskeleton Kit, tebu-bio; 1µM final concentration in liquid F12 culture medium) during 1 hour at 38°C, they were then mounted for en-face culture in LMP agarose medium, and incubated in SiR-tubulin at 100nM final concentration in liquid F12 culture medium.

### Compaction index measurement

The Compaction index (Ci) was determined using ImageJ software. Ci quantifies the effect of CatchFire pairing on mitochondrial compaction in dividing progenitors by measuring the ratio of a z-projection of the whole mitochondrial volume, imaged from the apical surface, to the surface of the maximum cell contour at the time cytokinesis (fig.S6D). The maximum cell contour of the pair of sister cells was manually outlined from a 3µm z projection of the equator region where the daughter cell pair’s surface area is maximum. Then a z projection of the total mitochondrial fluorescence from the pair of cells was performed, and contours of this fluorescence were manually drawn. For each pair of daughter cells, the ratio of the area occupied by mitochondria to the maximal projected area of the daughter cell pairs was calculated (fig.S6D). All control progenitors in the absence of ligand display ratios above 0.6 (fig.S6E). Accordingly, cells with a Ci value below 0.6 were considered as having compacted mitochondria.

### Immunostaining on vibratome sections

For vibratome sections, chick embryos were electroporated at E1.5 and harvested 30 hours later in ice-cold PBS. Extraembryonic membranes were quickly removed and embryos were fixed for 1 hour in ice-cold 4%PFA/PBS and rinsed 3 times for 5 minutes in PBS at room temperature (RT). The trunk region was then dissected and embedded in 4% agarose (4g of agarose in 100ml of water, boiled in a microwave and cooled at 50°C). 100 μm vibratome sections were obtained using a Microm HM 650 V Microtome (ThermoScientific) and collected in 6-well plates in ice-cold PBS.

Vibratome sections were incubated for one day with the primary antibodies diluted in PBS/0.1%Triton (PBST) at 4°C with gentle agitation. Sections were then washed 3 times for 5 minutes in PBST at room temperature (RT), incubated overnight in the dark at 4°C with the appropriate secondary antibodies and DAPI diluted in PBST, washed again 3 times for 5 minutes at RT with PBST and mounted with Vectashield (Vector Laboratories H-1000-10). Primary antibodies used are: rabbit anti-pRb (Ser807/811 – 1:1000) from Cell Signaling; mouse anti-HuC/D (clone 16A11 – 1:50) from ThermoFisher Scientific; Chick anti-GFP (1/800) from Aves Labs. Alexa Fluor 488-coupled anti-chicken (1:500) and Cy5-coupled anti-mouse (1:500) and anti-rabbit (1:250) secondary antibodies were obtained from Jackson laboratories.

## Key Resources

| REAGENT OR RESSOURCE | SOURCE | IDENTIFIER |
| --- | --- | --- |
| **Antibodies** |  |  |
| Chick anti-GFP | Aves Labs | Cat#GFP-1020<br>RRID : AB_10000240 |
| Rabbit anti-pRb (Ser807/811) | Cell Signaling | Cat# 8516S<br>RRID : AB_331472 |
| Mouse anti-HuC/D (clone 16A11) | ThermoFisher Scientific | Cat#A-21271<br>RRID : AB_221448 |
| **Softwares** |  |  |
| ImageJ | (35) | http://imagej.net/Welcome<br>RRID: SCR_003070 |
| Graphpad Prism | Graphpad | http://www.graphpad.com/<br>RRID: SCR_002798 |
| Microsoft Excel | Microsoft | RRID: SCR_016137 |
| MicroManager | (32) | https://micro-manager.org/<br>RRID: SCR_000415 |
| Affinity Publisher 2.5 | Affinity | N/A |
| IMARIS 10.0.0 | BitPlane | N/A |

---

**Table S1:**
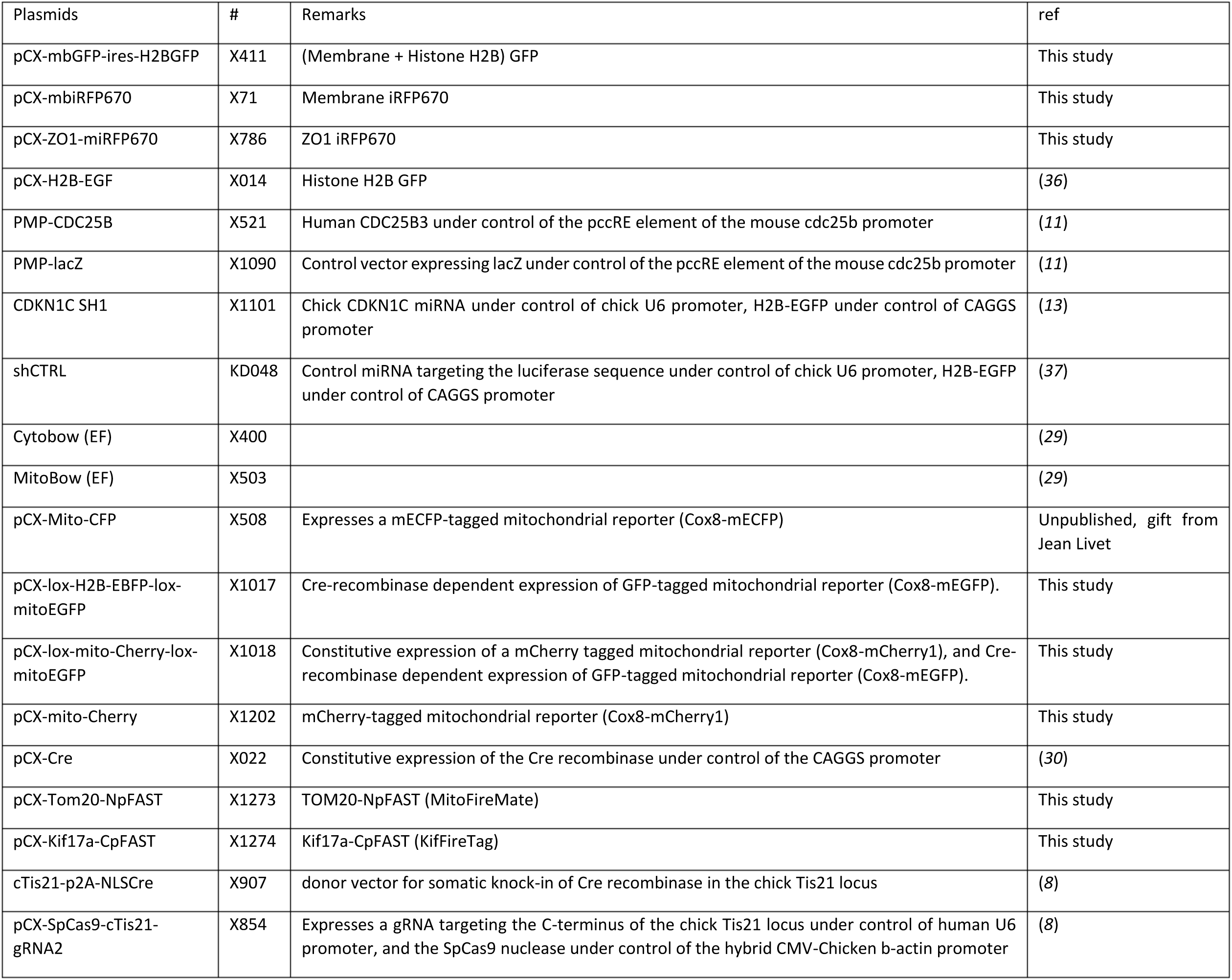
List of plasmids used in this study.

**Supplementary Figure 1:**
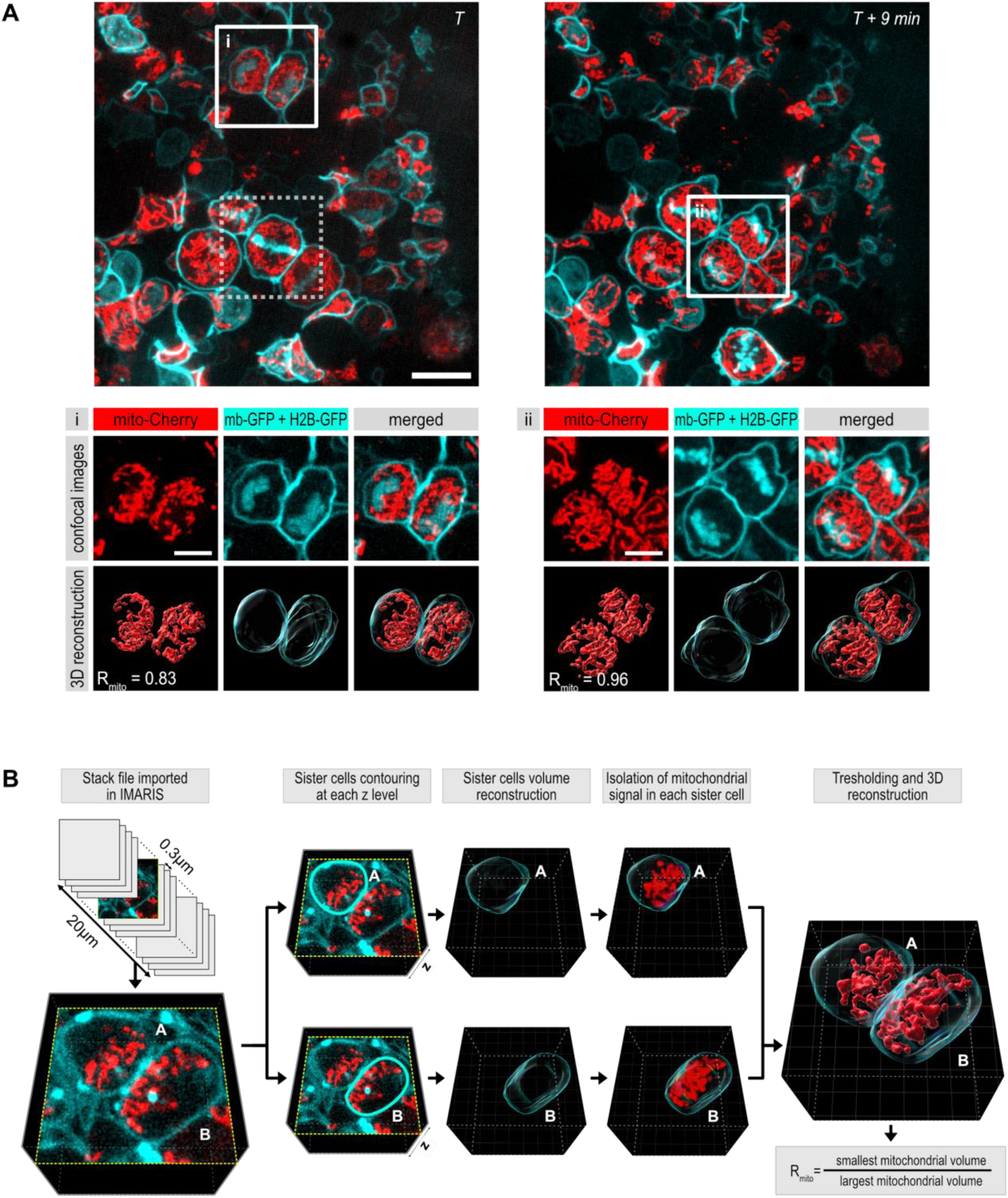
3D reconstruction of mitochondrial volume in daughter cells from a neural progenitor. Related to Figure 1. A. Top panels: en-face view of a single z-plane from a 90*90µm field of a neural tube in live imaging at two different time points. Red : Mitochondria, Cyan : cell contour. Bottom panels show images and 3D reconstructions of mitochondria in pairs of daughter cells of two progenitors dividing at successive timepoints at E2. The cells in i) and ii) correspond to white boxes in upper left and upper right panels, respectively). The dashed white box on the upper left panel highlights a metaphase progenitor whose daughter cells are visible in panel ii.. Upper rows in i) and ii): z plane of cell contour and chromosomes (middle, cyan), mitochondria (left, red), and merged (right) live fluorescence; Bottom rows: corresponding 3D reconstructed images from the entire z-stacks. Scale bars 5µm. B. **3D reconstruction pipeline:** Stack files of images of daughter cells immediately after division (∼25µm deep z-stacks with a 0.3µm z-step) were imported in IMARIS software (Bitplane) to perform cell volume and mitochondrial volume 3D reconstructions. Each daughter cell volume was delineated manually by drawing cell contour on each z plane through their entire height using either the membrane reporter signal or a cytoplasmic reporter signal, and reconstructed using the surface creation tool of IMARIS. The mitochondrial signal contained in the cellular volume of each daughter cell was then isolated by masking mitochondrial signal located outside of this volume. Each daughter’s mitochondrial volume was then reconstructed with the surface creation tool, using the same threshold for both cells. Values of mitochondrial volumes for each daughter cells were then used to calculate R_mito_.

**Supplementary Figure 2:**
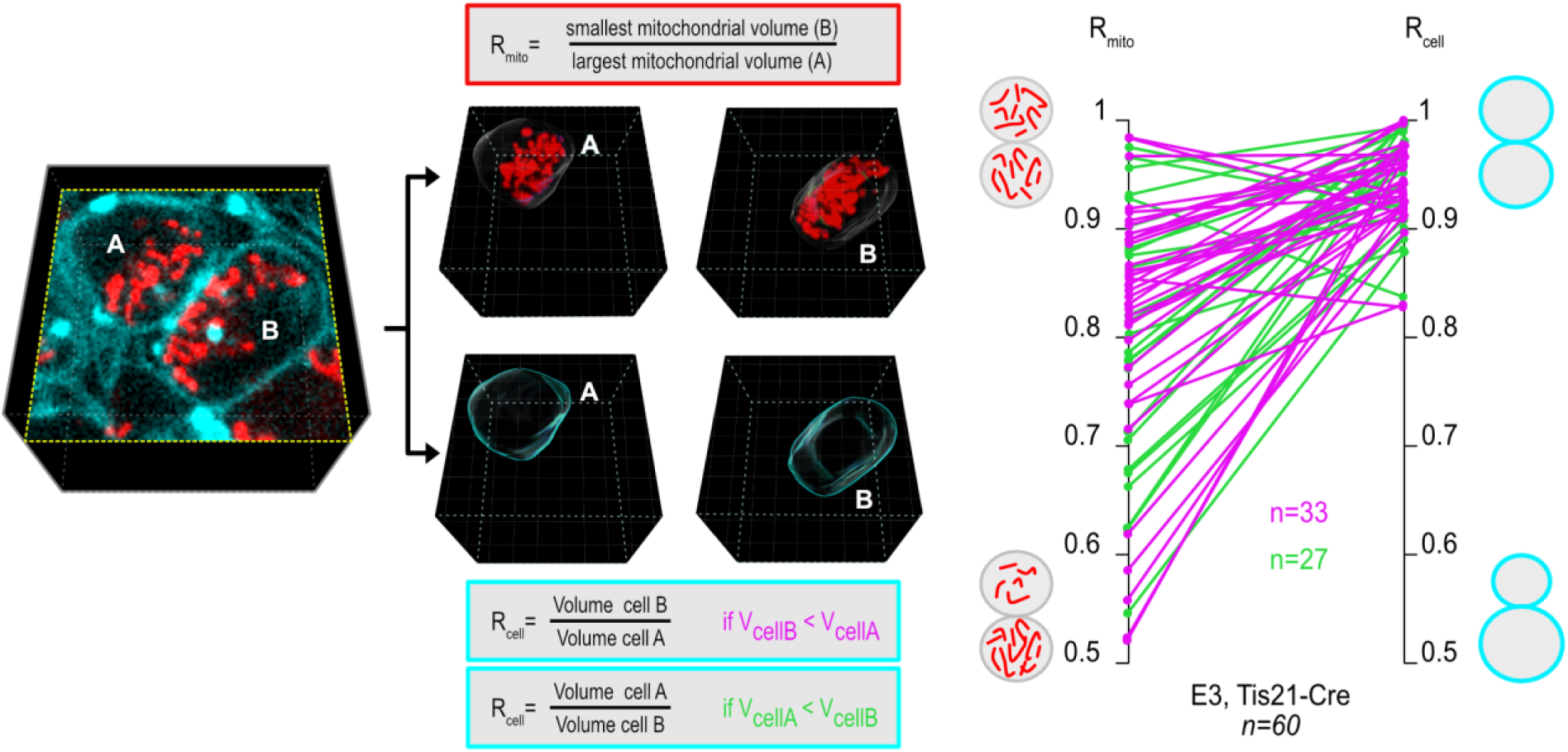
Absence of correspondence between mitochondrial inheritance ratio and cell volume ratio. Related to Figure 1. **Left:** 3D reconstruction of cell volume in pairs of daughter cells born from a neural progenitor. Stack file of images of daughter cells immediately after division (∼25µm deep z-stacks with a 0.3µm z-step) were imported in IMARIS software (Bitplane) to perform cell volume and mitochondrial volume 3D reconstructions. Each daughter cell volume was delineated manually by drawing cell contour on each z plane through their entire height using either the membrane reporter signal and reconstructed using the surface creation tool of IMARIS. The mitochondrial signal contained in the cellular volume of each daughter cell was then isolated by masking mitochondrial signal located outside of this volume. Each daughter’s mitochondrial volume was then reconstructed with the surface creation tool, using the same threshold for both cells. Data of both mitochondrial and cellular volumes for each daughter cells were then exported from Imaris to perform ratio measurements to assess the mitochondrial distribution between daughter cells and to compare their cellular volume. **Right:** Ratios of mitochondria volumes (R_mito_) are plotted on the left line. The corresponding cell volume ratios (R_cell_) are plotted on the right line. Magenta marks pairs in which the cell inheriting the smallest mitochondrial volume (cell B) is smallest than its sibling (cell A). Green marks pairs in which the cell inheriting the smallest mitochondrial volume (cell B) is larger than its sibling; in this case, R_cell_ corresponds to the ratio between the volume of cell A over the volume of cell B. N=60 pairs from Tis21-Cre expressing cells.

**Supplementary Figure 3:**
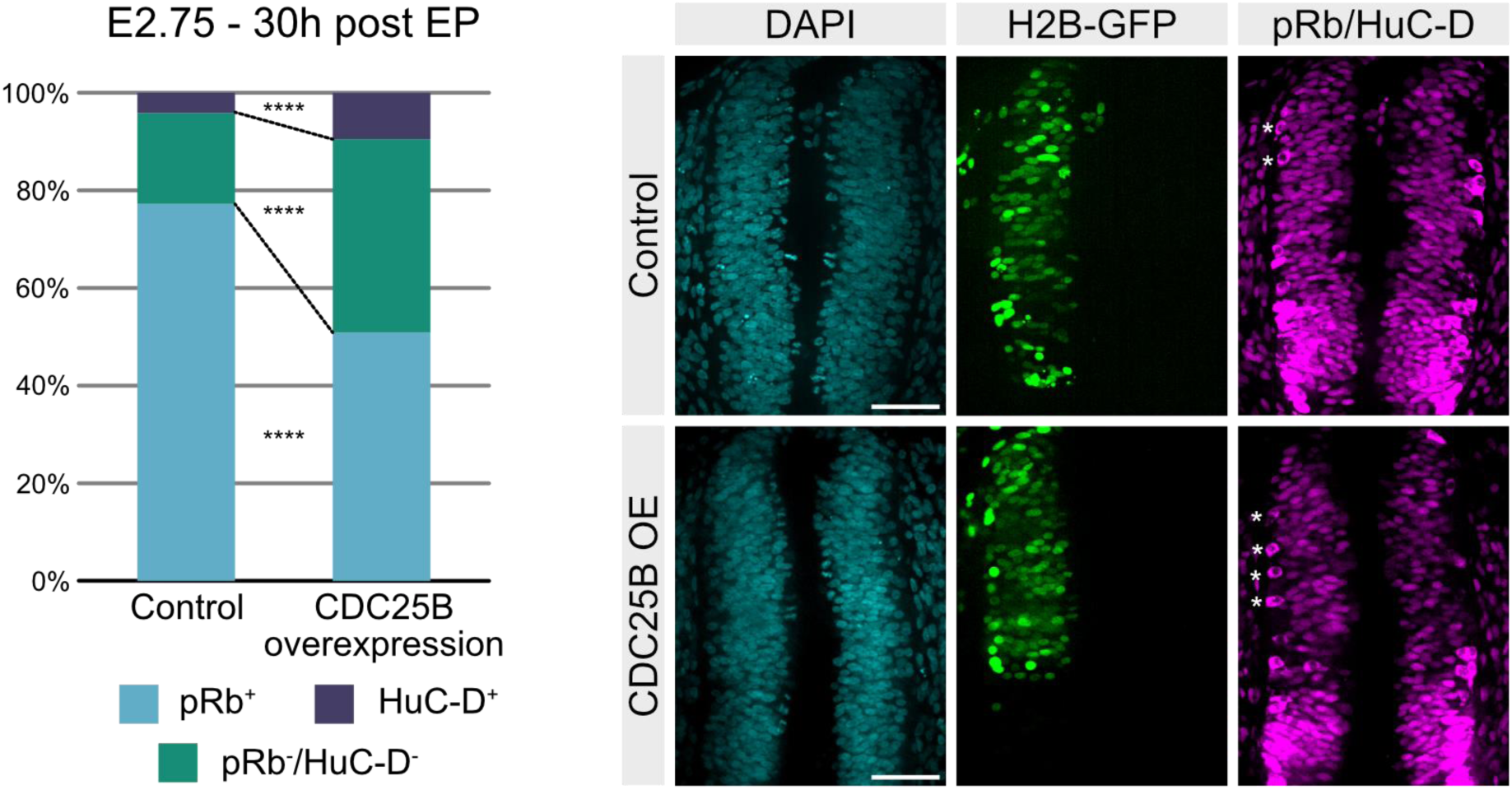
Increased neurogenesis upon Cdc25B overexpression in the chick embryonic neural tube. Related to Figure 2. **Left:** Distribution of the pRb positive cells (light blue), pRb/HuC-D double negative cells (green) and HuC-D positive neurons (dark blue) in control or CDC25B overexpression conditions at E3, 30 hours after electroporation (30 hours post EP). N=4 embryos for control and N=3 embryos for CDC25B overexpression conditions. Statistical analysis was performed using Mann-Whitney test ****: p < 0.0001. **Right:** Transverse sections of the electroporated chick neural tube (thoracic level) at E3. Immunofluorescence with HuC-D antibody to label neurons (magenta, cytoplasmic signal) and pRb to label progenitors (magenta, nuclear signal) (right panels) in control and overexpression of CDC25B (CDC25B OE) conditions. H2B-GFP fluorescence (middle panels, green) represents the fluorescent electroporation reporter and DAPI staining is showed in blue (left panels). Scale bars: 50µm

**Supplementary Figure 4:**
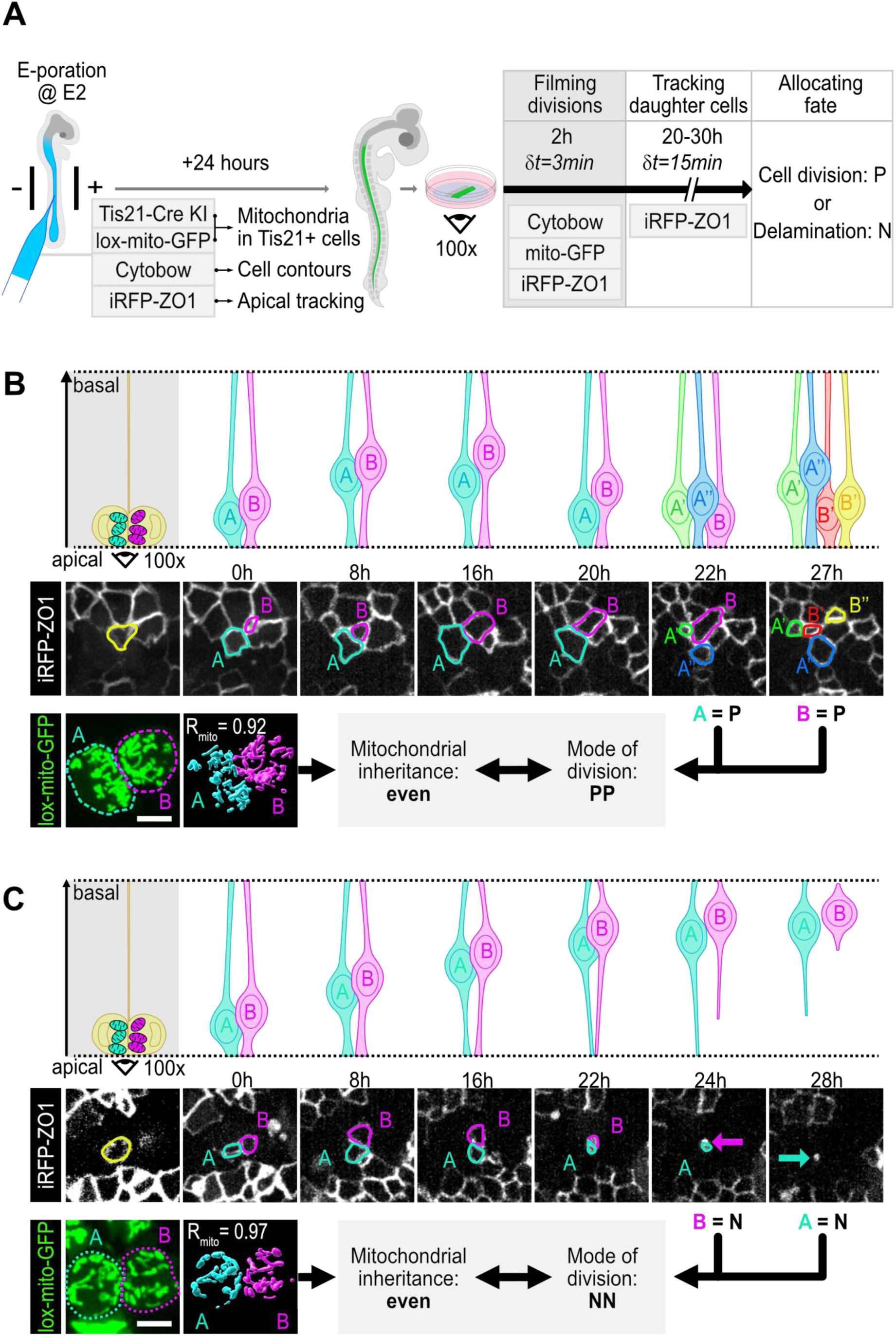
Fate tracking in pairs of daughter cells from a Tis21 positive progenitor showing even mitochondrial distribution and symmetric modes of division. Related to Figure 2. **A. Experimental strategy to combine live monitoring of mitotic mitochondrial inheritance and tracking of daughter cell fate.** *In ovo* electroporation of a combination of five vectors carrying DNA constructs for Tis21-P2A-Cre somatic knock-in, and fluorescent reporters for mitochondria (lox-mito-GFP), apical tracking (iRFP-ZO1) and cytoplasm staining (Cytobow) in the chick embryonic neural tube at E2. Embryos were harvested at E3, and neural tubes were mounted for en-face imaging. The neuroepithelium was imaged at high spatio-temporal resolution (δt=3min and z=0.3µm) for two hours to record mitochondrial inheritance during progenitor divisions, followed by tracking of the daughter cells by imaging the apical surface at lower resolution (δt=15min and z=1µm, 20µm stack height centered on iRFP-ZO1 staining) for 20 to 30 hours. B-C. Time-lapse series (en-face imaging) of dividing neural progenitors showing even mitochondrial distribution and symmetric PP (B) or NN (C) modes of division Top rows: Schematic representation of the time course from the division of a progenitor to the determination of its daughter’s fate based on their behavior and morphological criteria (new division for a progenitor, delamination for a neuron). Middle rows show time-lapse series (en-face imaging) of the iRFP-ZO1 signal used for long term tracking of pairs of daughter cells. In panel B, both daughter cells A and B are progenitors (as deduced from their division producing A’ and A’’, and B’ and B’’), while in panel C, both daughters are neurons (as deduced from the progressive reduction of their apical surface followed by delamination at 24 hours and 28 hours). Bottom rows: mitochondrial distribution (R_mito_) is measured immediately after cytokinesis of the mother cell and matched with the mode of division deduced from the fate of the daughters. Scale bars in B-D: 5µm.

**Supplementary Figure 5:**
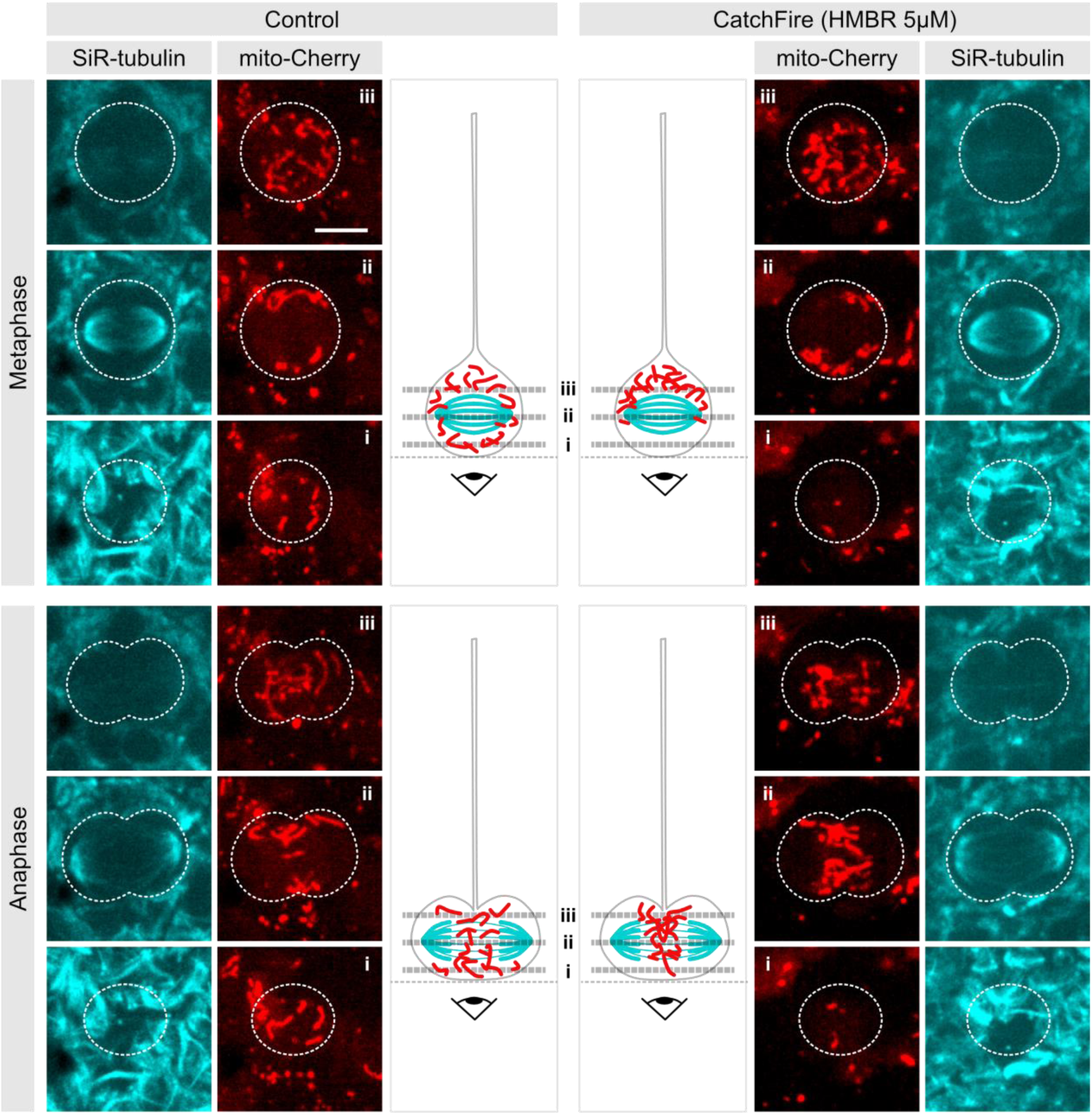
Mitochondrial behavior and localization during metaphase and anaphase are modified in CatchFire electroporated cells upon ligand (HMBR) administration. Related to Figure 3. z-projections (2.5µm corresponding to 5 z-levels) at three different levels (top, middle and bottom) of dividing progenitors, (see schematic representation in the middle panels) during metaphase, (upper panels) and anaphase (lower panels). Mitochondria (mito-Cherry, red), which are typically dispersed throughout the cytoplasm of the mitotic cell in metaphase in control conditions (left panels), are concentrated more basally in ligand exposure conditions (right panels). In anaphase, mitochondria appeared more compacted at the cleavage furrow upon ligand administration (right panels) compared to the control situation (left panels). Scale bars: 5µm

**Supplementary Figure 6:**
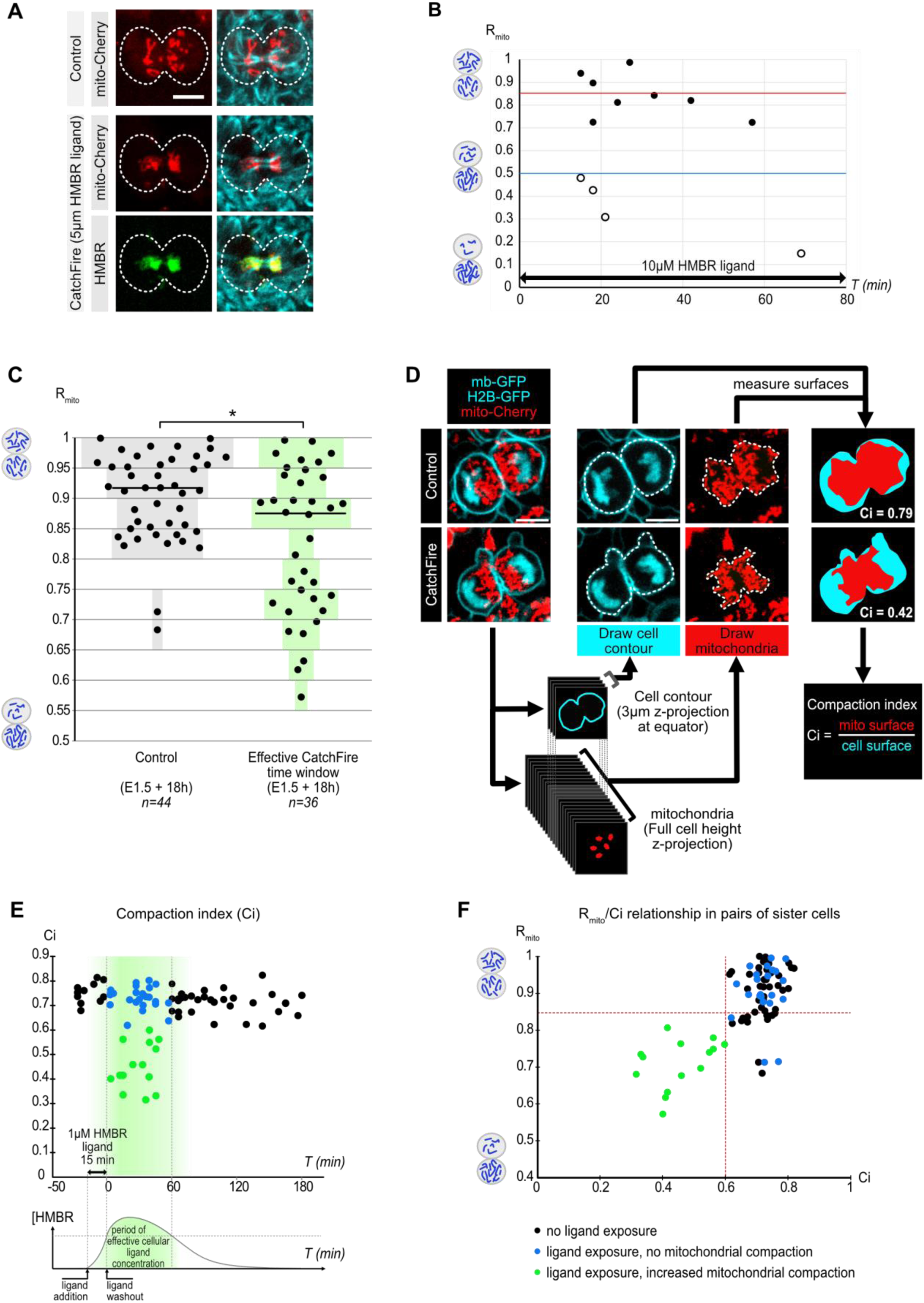
CatchFire-induced mitochondria/Kif17 interactions modify mitochondrial behavior and localization. Related to Figure 3. A. **Visualization of the sites of mitochondria/Kif17 interaction in telophase.** The interaction between the two partners (Tom20-FireMate, Kif17-FireTag) in the presence of the HMBR ligand is visualized via green fluorescence (HMBR, green) (see Figure 3A), which colocalizes with an independent mitochondrial reporter (mito-Cherry, yellow). The signal also partially colocalizes with the microtubular network (SiR-tubulin, cyan). Mitochondria are more compacted at the cleavage furrow in ligand exposure conditions (middle and lower panels) compared to control condition (upper panels). Scale bars: 5µm. B. **Graphical representation of mitochondrial ratios over time during continuous exposure to the HMBR ligand at high concentration (10µM).** Each dot represents a mitochondrial ratio. The red line shows the threshold (0.85) between “equal” and “unequal” R_mito_ values and the blue line shows the lower limit (0.5) of R_mito_ observed in control conditions. Open circles show cases of R_mito_ lower than 0.5, indicating an extreme imbalance in mitochondrial inheritance that is never observed in control conditions. The ligand was added to the culture medium at T=0min and maintained in the culture during the entire experimental period. C. **Scatter plots of R_mito_ in CatchFire condition before and after (Control) and during the estimated 1 hour period of effective intracellular CatchFire ligand concentration** defined in Figure 3B. Gray and green color bars represent the percentage of cell pairs in each 0.05 interval. Data are the same as in Figure 3B. Statistical analysis: Mann-Whitney. *: p=0.0137 n=80 pairs. D. **Schematic representation of the measurement of the Compaction index (Ci),** illustrated by live imaging of dividing progenitors (upper panels). To determine Ci, we used a z-projection of the total mitochondrial signal from the whole cellular volume (mito-Cherry, red) and a z-projection of the maximum cell contour (3 to 5 z-levels at the cell’s equator; mb-GFP, green) at the time of cytokinesis. The contours of the projected cell and mitochondrial surfaces were manually drawn (middle panel, green and red respectively). Ci is calculated as the ratio of the corresponding surfaces (mitochondrial over equatorial cell surface of the daughter cell pairs). Examples of control (top) and HMBR treated (bottom) sister cells are shown. Scale bars: 5µm. E. **Graphical representation of Ci in cells electroporated with the CatchFire components**, measured before (-50 to -15 min) addition of the ligand_,_ in presence of ligand in the medium (- 15 to 0 min), and following ligand removal from the medium (0 to 180 min). Each dot represents Ci for 1 pair of sister cells. Many cases of Ci values lower than 0.6 (green dots) are observed during a 1 hour time window after ligand removal from the medium (green box), whereas all values are higher than 0.6 before and after this time window. This corresponds to the time period during which we observed low values of R_mito_ in the same experiment (also marked with a green box in Figure 3B). Blue dots represent pairs measured during the period of effective ligand activity which retained a Ci similar to the control condition. Bottom: Scheme of the theoretical variation of the intracellular ligand concentration during the experimental period. Data are from the same dataset as in panel C and in Figure 3B, except that Ci could not be measured in one control pair. N=79 pairs. F. **Graphical representation of the relationship between R_mito_ and Ci values measured at E2.25** in cells electroporated at E1.5 with the CatchFire components, corresponding to the experimental dataset depicted in panels C and E and in Figure 3B. Blue and green dots correspond to cells dividing during the 1 hour time window highlighted by the green box in panel E and Figure 3B. Note that in this population, all the green cells with a Ci value lower than 0.6 (which is never observed in the control population) display a R_mito_ lower than 0.85, whereas most of the blue dots have similar R_mito_ as the control population. N=79 pairs.

**Supplementary Figure 7:**
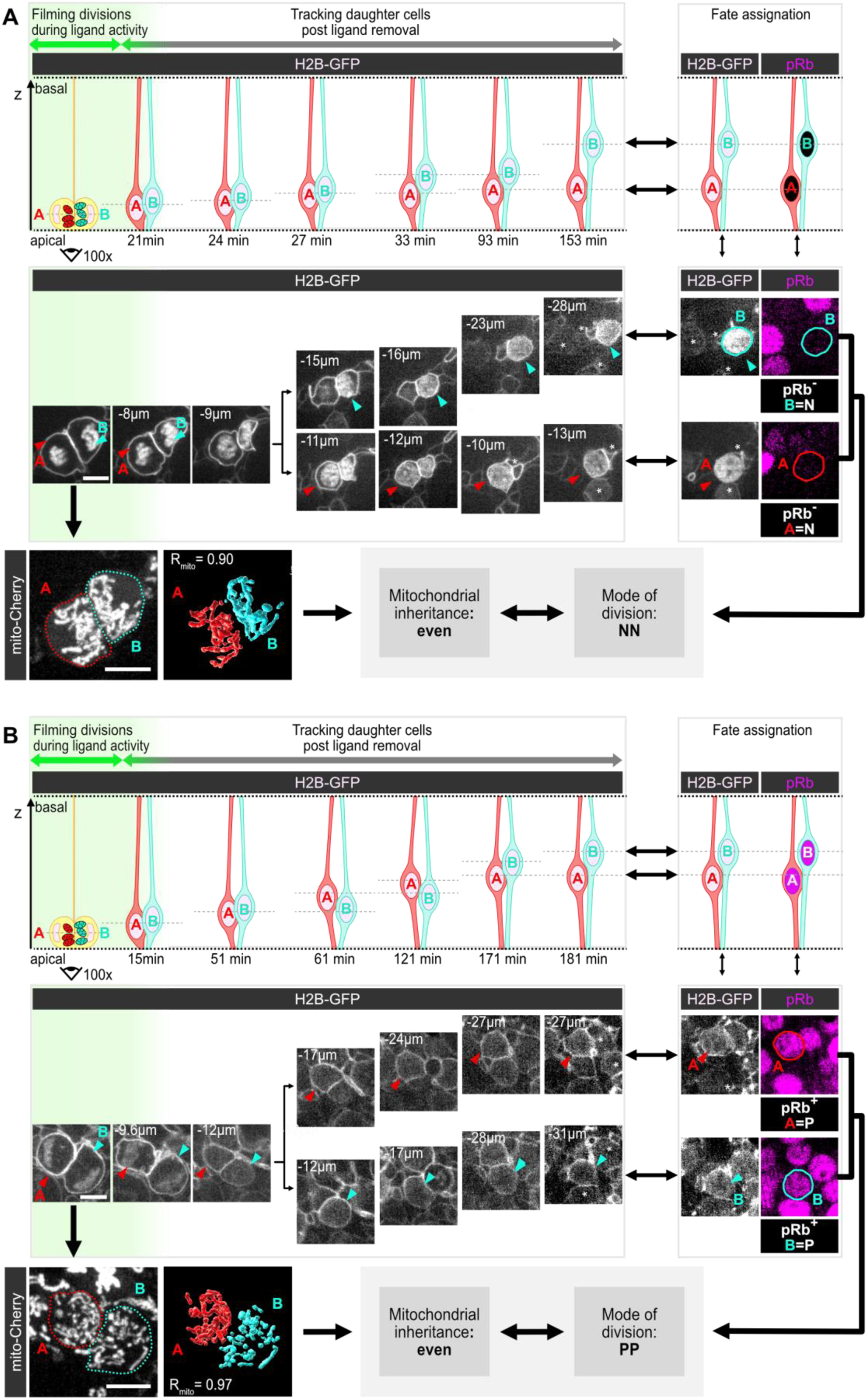
Forced unequal mitochondrial inheritance induces asymmetric fate choices. Related to Figure 3. Representative examples of a NN division **(A)** and a PP division **(B)** both displaying a characteristic equal mitochondrial inheritance. Scheme (top) and confocal en-face views (middle) of daughter cell’s nucleus (H2B-GFP, white) in the depth of the neuroepithelium during the live tracking period (left) and correspondence with pRb immunofluorescence at its end (right, magenta). Bottom: R_mito_ value measured upon cytokinesis of the mother cell is matched with daughter cells fate deduced from tracking: Scale bar: 5µm.

**Supplementary Figure 8:**
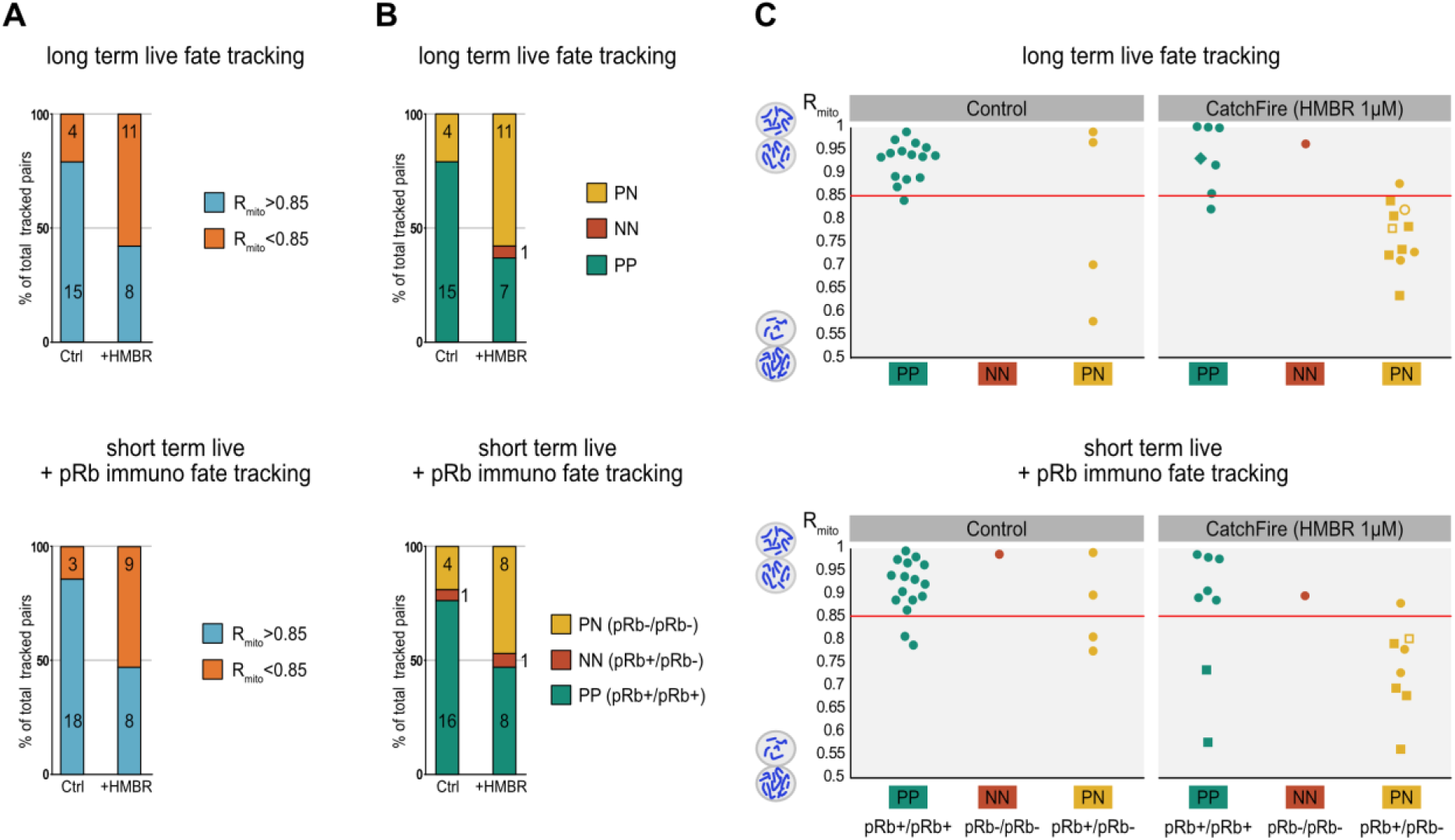
Forced unequal mitochondrial segregation in progenitors dividing mostly symmetrically leads to differential cell fate in daughter cells. Related to Figure 3. A. **Graphical representation of percentages of divisions displaying R_mito_ values higher versus lower than 0.85 in the live tracking experiments** in control and CatchFire conditions, from the long-term live tracking (top) and from the combined short tracking and immunostaining (bottom) experiments. Numbers indicated in columns correspond to cell numbers. Data are from the same experiments presented in a combined format in Figure 3E. B. **Graphical representation of percentages of PP, PN and NN pairs of sister cells** in control and CatchFire conditions in the live tracking experiments, from the long-term live tracking (top) and from the combined short tracking and immunostaining (bottom) experiments. Numbers indicated in columns correspond to cell numbers. Data are from the same experiments presented in a combined format in Figure 3F. C. **Correspondence between mitochondrial inheritance and fate of daughter cells** in control and CatchFire conditions. The datasets obtained from the two independent methods for fate assignation based either on long term live tracking (n=19 control cells and n=19 cells in CatchFire condition) or on combined short tracking and immunostaining (n=21 control cells and n=17 cells in CatchFire condition) are presented in the upper and lower rows respectively. They are from the same experiments presented in a combined format in Figure 3G.

## Legends to Supplementary Movies

Supplementary Movie 1: En-face time lapse monitoring of mitochondrial segregation during progenitor cellular divisions in chick embryonic neuroepithelium. Related to **Figure 1**.

Plasmids encoding mito-Cherry (red) and mb-iRFP (cyan) were electroporated in the neural tube *in ovo*, at embryonic day 2 (E2, HH stage 13–14). 24 hours later, embryos were dissected, and the neuroepithelium was imaged in en-face view every 3min during one hour. The movie displays a single focal plane from a z-stack, 5µm below the apical surface. White rectangles indicate pairs of sister cells immediately after cytokinesis, corresponding to the time points used for 3D reconstructions and measurements of mitochondrial volume inherited by each sister in this study. Scale bar, 20µm.

Supplementary Movie 2. Mitochondrial segregation during cellular division of a neural progenitor in the chick embryonic neuroepithelium. Related to Figure 1 and Supplementary Figure 1.

Plasmids encoding mito-Cherry (red) and a mb-GFP-ires-H2B-GFP (cyan) were electroporated in the neural tube *in ovo*, at embryonic day 2 (E2, HH stage 13–14) and the neuroepithelium was imaged in en-face view 24 hours later. Upper row of the movie: single focal planes at 4 consecutive timepoints from metaphase to cytokinesis of a single progenitor, 6 µm below the apical surface. Bottom row shows sequential views of all single z-planes within the z-stack in metaphase (left) and after cytokinesis (right). The latter was imported in Imaris for 3D- reconstruction of mitochondrial and cellular volumes in the two daughter cells (middle). In this example, both daughter cells inherit similar mitochondrial volumes from the mother cell (R_mito_=0.97). Scale bar in confocal images, 5µm.

Supplementary Movie 3. Combined monitoring of mitochondrial inheritance and sister cell fate in an asymmetrically dividing progenitor (PN). Related to Figure 2.

Plasmids encoding fluorescent reporters for mitochondria (lox-mito-GFP), apical tracking (iRFP-ZO1), cytoplasm staining (Cytobow, used to identify the cellular contour for 3D reconstruction; not shown in the movie) and constructs for the targeting of the Cre recombinase at the Tis21 locus were electroporated in the neural tube at embryonic day 2 and the neuroepithelium was imaged in en-face view 24 hours later. The left panel shows the 3D reconstruction of mitochondrial volumes inherited by each sister A and B, indicating unbalanced inheritance (R_mito_ = 0.61) with B receiving a smaller volume. The right panel shows a time-lapse series of the apical iRFP-ZO1 signal used for long term tracking of pairs of daughter cells. Daughter cell B undergoes a progressive reduction of its apical surface followed by delamination at 24 hours and is therefore identified as a neuron, while daughter cell A divides at 26 hours and is therefore identified as a progenitor. Note that the daughter cell inheriting the smallest mitochondrial volume becomes a neuron.

Supplementary Movie 4. Combined monitoring of mitochondrial inheritance and sister cell fate in a symmetric proliferative progenitor division (PP). Related to Figure 2 and Supplementary Figure 4.

Plasmids encoding fluorescent reporters for mitochondria (lox-mito-GFP), apical tracking (iRFP-ZO1) and cytoplasm staining (Cytobow, used to identify the cellular contour for 3D reconstruction; not shown in the movie) and constructs for the targeting of the Cre recombinase at the Tis21 locus were electroporated in the neural tube at embryonic day 2 and the neuroepithelium was imaged in en-face view 24 hours later. The left panel shows the 3D reconstruction of mitochondrial volumes inherited by each sister A and B, indicating balanced inheritance (R_mito_ = 0.92). The right panel shows a time-lapse series of the apical iRFP-ZO1 signal used for long term tracking of pairs of daughter cells. Both daughter cells undergo a new round of division at 22 hours (A) and 26 hours (B) and are therefore identified as progenitors.

Supplementary Movie 5. Combined monitoring of mitochondrial inheritance and sister cell fate in a symmetric neurogenic progenitor division (NN). Related to Figure 2 and Supplementary Figure 4.

Plasmids encoding fluorescent reporters for mitochondria (lox-mito-GFP), apical tracking (iRFP-ZO1) and cytoplasm staining (Cytobow, used to identify the cellular contour for 3D reconstruction; not shown in the movie) and constructs for the targeting of the Cre recombinase at the Tis21 locus were electroporated in the neural tube at embryonic day 2 and the neuroepithelium was imaged in en-face view 24 hours later. The left panel shows the 3D reconstruction of mitochondrial volumes inherited by each sister A and B, indicating balanced inheritance (R_mito_ = 0.97). The right panel shows a time-lapse series of the apical iRFP-ZO1 signal used for long term tracking of pairs of daughter cells. Both daughter cells undergo a progressive reduction of their apical surface followed by delamination at 24 hours (B) and 30 hours (A) and are therefore identified as neurons.

Supplementary Movie 6. Identification of a “PN” pair of sister cells using a combination of live tracking after cytokinesis and pRb immunofluorescence. Related to Figure 3.

Sister cells expressing mb-GFP and H2B-GFP, CatchFire components and a mitochondrial reporter (mito-Cherry) were imaged in en-face cultures of the neuroepithelium at E2.25. z-stacks were acquired at 3 min intervals and 0.3µm z steps for 1 hour, then at 15 min intervals and 1µm z steps for an additional 2.5 hours. The two left columns show single focal planes from the live sequence. Only the GFP channel is shown. Due to interkinetic nuclear movements, both sisters’ nuclei move in the depth of the neuroepithelium at different speeds, and are therefore displayed on two different rows (top and bottom) from 45 min onwards until the end of the live imaging sequence. Samples were fixed at the end of the live imaging sequence and processed for immunostaining with anti-GFP and pRb antibodies. The third column shows the correspondence of the anti-GFP signal in the fixed sample with the last time point of the live tracking sequence, and the 4^th^ column shows the pRb signal. Cell A (top row) is pRb negative, and therefore identified as a neuron, while cell B (bottom row) is pRb positive and identified as a cycling progenitor. T=0 min corresponds to the timing of cytokinesis of the mother cell. Asterisks indicate neighboring cells used for registration between live and fixed images.

Supplementary Movie 7. Identification of a “PP” pair of sister cells using a combination of live tracking after cytokinesis and pRb immunofluorescence. Related to Figure 3 and Supplementary Figure 7.

Sister cells expressing mb-GFP and H2B-GFP, CatchFire components and a mitochondrial reporter (mito-Cherry) were imaged in en-face cultures of the neuroepithelium at E2.25. z-stacks were acquired at 3 min intervals and 0.3µm z steps for 1 hour, then at 10 min intervals and 1µm z steps for an additional 2 hours. The two left columns show single focal planes from the live sequence. Only the GFP channel is shown. Due to interkinetic nuclear movements, both sisters’ nuclei move in the depth of the neuroepithelium at different speeds, and are therefore displayed on two different rows (top and bottom) from 49 min onwards until the end of the live imaging sequence. Samples were fixed at the end of the live imaging sequence and processed for immunostaining with anti-GFP and pRb antibodies. The third column shows the correspondence of the anti-GFP signal in the fixed sample with the last time point of the live tracking sequence, and the 4^th^ column shows the pRb signal. Both cell A (top row) and cell B (bottom row) are pRb positive and identified as cycling progenitors. T=0 min corresponds to the timing of cytokinesis of the mother cell. Asterisks indicate neighboring cells used for registration between live and fixed images.

Supplementary Movie 8. Identification of a “NN” pair of sister cells using a combination of live tracking after cytokinesis and pRb immunofluorescence. Related to Figure 3 and Supplementary Figure 7.

Sister cells expressing mb-GFP and H2B-GFP, CatchFire components and a mitochondrial reporter (mito-Cherry) were imaged in en-face cultures of the neuroepithelium at E2.25. z-stacks were acquired at 3 min intervals and 0.3µm z steps for 1 hour, then at 10 min intervals and 1µm z steps for an additional 2 hours. The two left columns show single focal planes from the live sequence. Only the GFP channel is shown. Due to interkinetic nuclear movements, both sisters’ nuclei move in the depth of the neuroepithelium at different speeds, and are therefore displayed on two different rows (top and bottom) from 45 min onwards until the end of the live imaging sequence. Samples were fixed at the end of the live imaging sequence and processed for immunostaining with anti-GFP and pRb antibodies. The third column shows the correspondence of the anti-GFP signal in the fixed sample with the last time point of the live tracking sequence, and the 4^th^ column shows the pRb signal. Both cell A (top row) and cell B (bottom row) are pRb negative, and therefore identified as neurons. T=0 min corresponds to the timing of cytokinesis of the mother cell. Asterisks indicate neighboring cells used for registration between live and fixed images.

